# Expanding the scope of a catalogue search to bioisosteric fragment merges using a graph database approach

**DOI:** 10.1101/2024.08.02.606367

**Authors:** Stephanie Wills, Ruben Sanchez-Garcia, Stephen D. Roughley, Andy Merritt, Roderick E. Hubbard, Frank von Delft, Charlotte M. Deane

## Abstract

The efficiency of fragment-to-lead optimization could be improved by automated workflows for the design of follow-up compounds. Pipelines that are able to fully exploit the interaction opportunities identified from the crystal structures of bound fragments would greatly aid this goal. To do so, these pipelines need to require minimal intervention from the user and be computationally efficient. In this work, we describe an updated version of our fragment merging methodology, which provides several feature enhancements, primarily by expanding the chemical space searched, allowing the identification of more diverse follow-up compounds, thus maximizing the chances of finding successful hits. While the original method focused on finding ‘perfect merges’, meaning compounds that directly incorporate substructures from the original fragments, here we expand the search to what we term ‘bioisosteric merges’, involving the incorporation of substructures that replicate the pharmacophoric features of the original fragments but may not be exactly identical. Unlike existing pharmacophore and shape-based descriptors used for virtual screening, this approach combines the search for these properties with the incorporation of novelty, which is necessary when searching for ways to link together distinct substructures. Compared with ‘perfect merging’, our new approach is able to find compounds that are directly informed by structures within the original fragments but are more chemically diverse. We contrast our approach with the use of a pharmacophore-constrained docking pipeline, run in parallel for select fragment pairs, and show that our method requires between 1.1-45.9-fold less computational time for conformer generation per merging ‘hit’ identified, referring to compounds that show a favourable degree of shape and colour overlap and recapitulation of original fragment interactions. Overall, our results show that our method has potential to be used to generate designs inspired by all fragments within a given pocket.

## 1 Introduction

Fragment-based drug discovery (FBDD) is now well-established as a technique to generate novel compounds that bind to and affect the activity of a protein of interest [1]. The approach starts with identifying small molecules (typically with less than 18 non-hydrogen atoms) that bind to the target using a sensitive screening technique such as NMR, surface plasmon resonance, a suitable biochemical assay or X-ray crystallography. Most of the successful FBDD projects have relied on protein-fragment crystal structures to then guide and control optimization of these small molecular starting points to become larger, more potent compounds. Recent developments in X-ray crystallographic screening and the associated data collection and structure determination have dramatically increased the number of fragment-bound crystal structures that can be obtained [2, 3, 4].

However, the design of follow-up compounds that exploit all possible interaction opportunities from a fragment screen in an automated manner is less routine. Existing pipelines have proved to be effective in progressing fragments to become follow-up compounds with substantially improved affinity (low ^M to nM), but have typically been restricted to scenarios with few (up to five) fragment hits [5, 6, 7]. There are several approaches for developing follow-up compounds based on crystal structures of bound fragments. Typically, these methods perform docking of virtual chemical libraries using template-guided docking based on the initial fragments. A major obstacle with docking tools is that they are computationally intensive, and thus docking entire compound libraries containing millions of compounds is not always practical. Consequently, appropriate prefiltering steps to select promising compounds are needed. This can be achieved, for instance, through the use of molecular fingerprint-based similarity searches [6] or the use of workflows that iteratively dock compounds using reaction-based enumeration [7]. Another issue is that docking results are far from perfect [8], hence many candidate compounds have to be tested in order to achieve a sufficient hit rate.

Our 2023 paper [5] provided an initial pipeline to address this problem. The original pipeline used a graph database representation of chemical space, termed the Fragment Network [9], to find catalogue compounds that contain substructures from two parent fragments simultaneously. The merges can then be passed through a set of 2D and 3D filters to prioritise compounds most likely to adopt the binding pose of the original fragments. Retrospective analysis of our merging pipeline was able to identify potential binders with low double-digit half-maximal inhibitory concentration (IC50) values.

In this new version of our algorithm, we have addressed some of its previous limitations, such as merging substructures that may not be spatially compatible or important for binding, or restricting our focus to ‘perfect merges’ — compounds that contain exact substructures of the parent fragments [5]. The original tool enumerated all pairs of fragments for merging, removing only pairs that are near identical in terms of chemical structure and overlap, and that exceed a specified distance threshold. Our new pipeline prioritises the merging of spatially compatible substructures that, optionally, make important interactions with the protein. Additionally, instead of limiting the search to exact substructures found in the initial fragment hits, it can find merges containing substructures that replicate pharmacophoric properties using pharmacophore fingerprints. For the updated algorithm, similarity metrics to determine appropriate replacement substructures are calculated on-the-fly as the database is queried.

This new search method combines both the power of similarity and substructure searches (with the Fragment Network representing a form of substructure search), and we refer to these newly identified types of compounds as ‘bioisosteric merges’. The term ‘bioisosteres’ refers to compounds where a functional group or fragment has been swapped with another that exhibits similar or enhanced biological activity. Tools for performing bioisosteric replacements have been developed to suggest substitutions for a given functional group based on various properties. For example, the Craig Plot [10] suggests bioisosteric replacements on the basis of hydrophobicity or electron donating and accepting power. Additionally, there are recommenders for bioisosteric ring [11] and linker replacements [12], which are designed to increase bioactivity. Our tool provides the ability to find bioisosteres within the context of finding fragment merges and linkers, requiring the introduction of novelty in how to join the bioisosteric replacements to create a physically reasonable compound that maintains the orientations of the inspirational fragments. There are many 3D shape and pharmacophore-based tools that have been developed to enable the identification of structural analogues for a given compound based on these properties. These tools are useful as they can enable us to perform virtual screening of large compound libraries by finding analogues for a known hit. Some examples of these algorithms include ultrafast shape recognition (USR) [13], which refers to a shape-based descriptor that is calculated from statistical moments using distances between reference atoms within the molecule. The USRCAT descriptor [14] builds on USR by considering the CREDO atom types [15], while ElectroShape [16] is another USR extension that considers electrostatic complementarity. Other methods utilise distributions based on distances between pharmacophores, such as Ligity [17], which calculates distances between triangular and tetrahedral pharmacophoric interaction points (interaction hot spots derived from protein-ligand complexes). To our knowledge, there is little evidence in the literature of the use of the above approaches within the context of fragment linking or merging. These descriptors are more commonly used for analogue searching, as linking and merging often require the generation of novelty. A method that approaches this is FRESCO [18], which uses unsupervised machine learning methods to learn 3D distributions of pharmacophores from a set of crystallographic fragment hits, which were in turn used to score and prospectively screen compounds from Enamine REAL space.

The potential benefits of expanding the search to bioisosteric merges include improved productivity and hit rates and maximizing the chemical and functional diversity of follow-up compound designs in the early stages of the FBDD pipeline. Maintaining this diversity is important as it increases the chances of developing promising leads further down the line (as chemically similar molecules are likely to exhibit similar biological properties [19]). Moreover, as a candidate hit is developed into a larger molecule, it becomes increasingly difficult for the chemist to make changes to the core scaffold. Our approach effectively combines elements of fragment optimisation, whereby the structure of a fragment hit is optimised within fragment space prior to elaboration, with the traditional fragment elaboration techniques of merging and linking. Thus, early exploration of chemical space is important, enabling a comprehensive understanding of the structure-activity relationship (SAR) and of which substructures are required for (or will ablate) binding [20, 21].

We find that this approach of expanding the search to bioisosteric merges enables a broader search of the chemical space represented in the Fragment Network, resulting in compounds that incorporate more unique substructures than when performing perfect merging alone. As a consequence, we identify merges that represent a greater number of fragment pairs and are able to access potential interactions that are not seen for the set of perfect merges. We use a conformer generation method that attempts to recapitulate the poses of the original fragments in a physically reasonable way. Our bioisosteric merges are predicted to adopt poses that indicate comparable shape and pharmacophoric overlap with the crystallographic fragments and thus represent a promising way of increasing diversity and improving the potential hit rate of follow-up compound design.

Comparison against a more traditional, ‘out-of-the-box’ docking approach using pharmacophoric constraints shows that our tool provides substantial benefits with regard to computational efficiency. We find, when comparing follow-up compounds proposed for specific fragment pairs, the time required for conformer generation to identify a computational merging ‘hit’ is 1.1-45.9-fold greater for the docking protocol. This suggests our method offers a practical and formulaic approach for enumerating follow-up compounds at scale using data from an entire crystallographic screen (rather than a few individual fragment pairs). A case study using retrospective experimental data demonstrates that our bioisosteric merging method mimics substructure variations that have been manually designed by chemists when exploring SAR.

## 2 Methods

### 2.1 Crystallographic fragment screening data

To test our merging pipelines, we used the results of XChem crystallographic fragment screens against three antiviral targets, the data for which can be freely downloaded from the Fragalysis platform (https://fragalysis.diamond.ac.uk/viewer/react/landing).

The first two targets are proteases from two non-polio enteroviruses (EVs), which are members of the Picornaviridae family, EV D68 and EV A71. EV D68 is most commonly associated with mild respiratory illnesses, although has been linked (in rare cases) to acute flaccid myelitis [22]. EV A71 is linked to hand, foot and mouth disease, which affects children, but can also cause severe neurological diseases, including encephalitis [23]. We looked specifically at the 3C protease (of EV D68) and the 2A protease (of EV A71). These proteases are involved in self-cleavage of the viral polypeptide into individual viral proteins and are also involved in the cleavage of several host cell proteins [22]. For EV D68 3C protease, we searched for merges for 25 hits found to bind to the catalytic site (Figure 1a). For EV A71 2A protease, we examined 6 hits that were found to bind to the P1 region of the active site (Figure 1b). The third target belongs to the Zika virus, which is a flavivirus that has been linked to neurological disorders, including Guillain-Barre syndrome, and the increased risk of microcephaly in fetuses and newborns owing to prenatal Zika infection [24]. The virus consists of an ^11kb RNA genome, which encodes a large polyprotein and is processed into several structural proteins and non-structural proteins. We focused on the NS2B-NS3 protease, which plays an essential role in processing the polyprotein to produce the virus’ enzyamtic components and the replication complex [25, 26]. The fragment screen shows a dense population of fragments in the S1 region of the active site for the NS2B co-factor, with 38 fragments found to bind, but providing several linking opportunities to one fragment in the S1‘ site and 3 fragments in the S2 site (Figure 1c). All fragments used in our analysis are listed in Supplementary Table 1.

**Figure 1:**
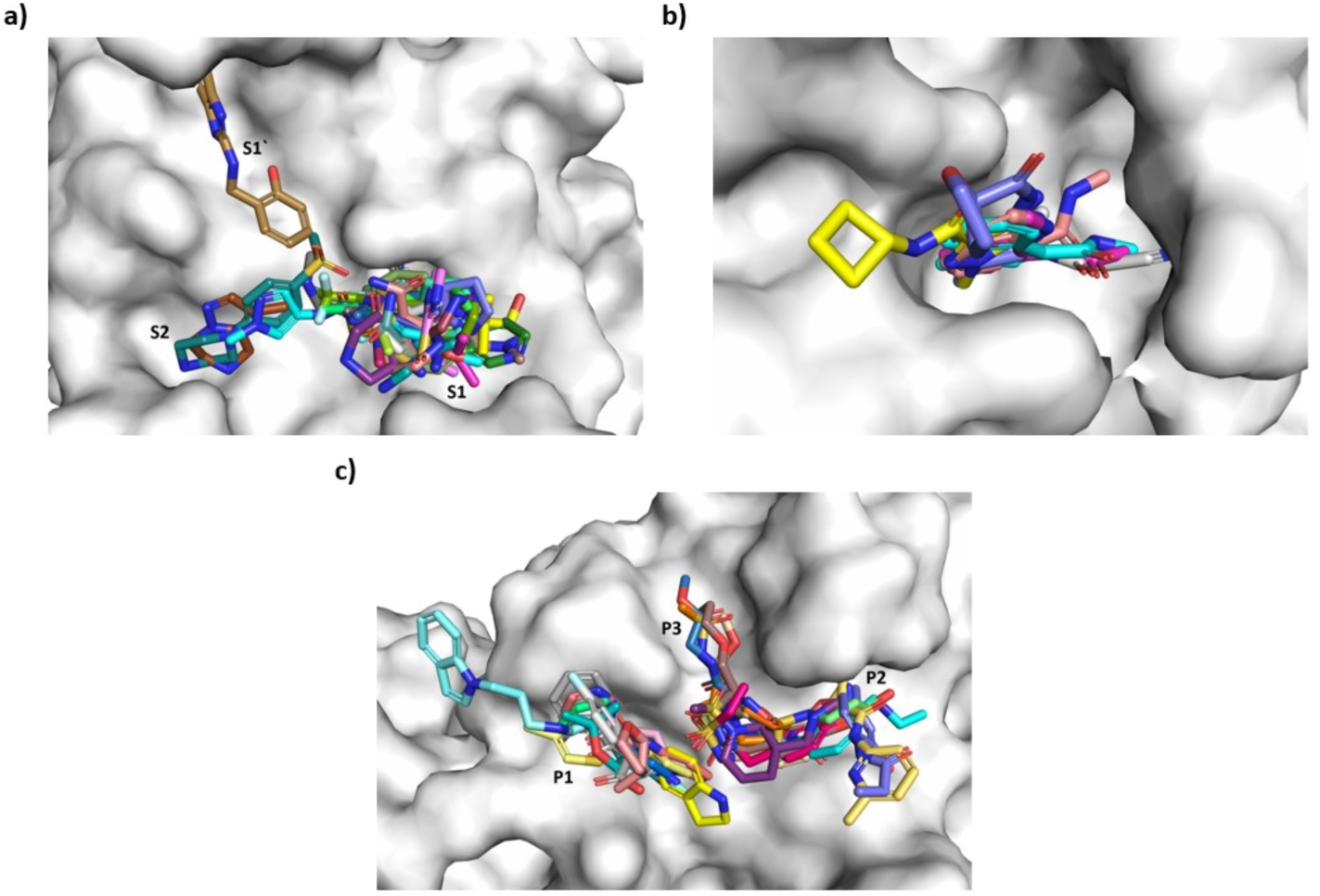
Crystallographic fragment hits used for analysis. The binding sites and crystallographic fragment hits used for testing the merging pipelines are shown for (**a**) enterovirus (EV) D68 3C protease, (**b**) EV A71 2A protease and (**c**) Zika NS2B. Individual subsites are labelled. Data can be freely downloaded from https://fragalysis.diamond.ac.uk/viewer/react/landing. The fragment codes are provided in the Supplementary Information.

### 2.2 Updates to the Fragment Network database

The Fragment Network version used in this work contains >200 million compounds from vendors, including Enamine, MolPort and Chemspace. It is an expansion of the version described in [5]. The network comprises nodes, representing molecules or substructures, which are connected by edges that denote transformations in which rings, linkers and substituents are added or removed between nodes. Purchasable compounds are labelled, enabling searches for vendor compounds resulting from specific types of transformation that are defined using the Cypher query language used by the neo4j graph database platform [27].

#### 2.2.1 Pharmacophore descriptors for substructures involved in transformations

We have added pharmacophore properties to the Fragment Network to enable the user to identify transformations involving substructures with desired pharmacophore properties. To do this, the edges of the Fragment Network are now additionally labelled with 2D fingerprints describing the pharmacophore features of the substructure being added or removed in a transformation. This allows the creation of queries that search for merge compounds that incorporate substructures with similar pharmacophore features to a query substructure (the full querying process is described below). In the context of this work, we focus on using 2D rather than 3D pharmacophore descriptors, as they have a reduced requirement for computational resources (as conformer generation is not needed) and we found that they were sufficient when searching for similar substructures. Comparison of 2D versus 3D fingerprints has found that the former perform as well as state-of-the-art 3D methods for multiple tasks, including toxicity, solubility and simple binding affinity prediction [28] (although their use is still possible in graph database searches).

The 2D pharmacophore fingerprint is calculated using the cheminformatics toolbox RDKit [29]. The fingerprint is generated by calculating topological distances between defined pharmacophore features, which are stored as bits in the fingerprint. We consider topological distances between pairs of pharmacophores, which are binned into three bins representing distances of 0—2, 2—5 and 5—8 bonds (to keep the length of the pharmacophore fingerprint short, resulting in fingerprints that were 84 bits in length). The pharmacophore features considered include hydrogen bond acceptors, hydrogen bond donors, negative ionizable, positive ionizable and aromatic groups. Two ‘pseudo-pharmacophore’ features were also added: aliphatic rings, as this was found to help maintain the shape of the substructure, and the attachment point atoms, which enables consideration of the location of the attachment point in relation to the rest of the substructure. Example substructure replacements are shown in Supplementary Figure 1.

### 2.3 Enumeration of substructures for querying

In the previous iteration of the Fragment Network merge search, the database was queried using pairs of fragments. In this new iteration, the search is performed for compatible pairs of substructures sourced from the fragments. Without controlling which final substructures are incorporated into the merge, the merge may occur between substructures that are not spatially compatible, wasting computational time.

First, as in our previous version of the tool [5], pairs of fragments are pre-filtered according to whether they share overlapping volume over a specified threshold, as these pairs do not represent good opportunities for merging (as they are unlikely to result in increased interactions). In this iteration of the tool, we removed pairs for which at least 56% of the volume of one fragment overlaps with the other. This threshold was determined by manually filtering pairs of fragments by visual inspection for an example fragment screen (and by calculating the threshold based on those that were filtered out). As in our previous iteration [5], we additionally removed fragment pairs that show a minimum distance between the closest pair of atoms of >5A. Both thresholds can be easily modified according to the needs of the user.

Following the enumeration of compatible fragment pairs, fragments are broken down by retrieving their constituent substructures from the network. Substructures are checked for whether they contain at least three carbon atoms (representing a substantial contribution to the final merge) and that they do not represent a single carbon-only ring (as these substructures are densely connected in the network and result in large numbers of molecules for filtering; this filter can be controlled by the user). To ensure the merges maximize binding, substructures are also (optionally) filtered for those that make an interaction with the protein (calculated using ProLIF [30]). We limited the number of compounds retrieved from the network, as some queries can result in large numbers of compounds for filtering. This is particularly common for ‘terminal nodes’ in the decomposition, meaning substructures that do not undergo any further contraction (such as single rings), which are extremely densely connected in the network. For each database query between substructures, results are limited to the first 500 compounds retrieved, although this limit is optional and can be easily modified. The parameters used are described in the Results section.

Substructures are filtered to remove those for which the geometric mean of the overlapping volume for each substructure (meaning the percentage volume of each substructure that is overlapping with the other) is ≥30%. The set of non-redundant substructure pairs (as the equivalent substructures can be found in multiple pairs of fragments) is subsequently used for querying the database. Example pairs of substructures for querying are shown in Supplementary Figure 2.

### 2.4 Querying the database

As the query is performed using substructures instead of fragments, the parameters of the search have been significantly modified compared with the previous iteration in [5]. The database query operates as follows: queries are performed for pairs of substructures (derived from two parent fragments) to be merged into a single compound. For clarity, we refer to the two substructures as substructure A and substructure B. Substructure A represents the ‘seed node’ in the query, and the query retrieves the database node representing this substructure to initiate the search. To identify perfect merges, from the seed node, up to two optional hops are made, in which an expansion can be made, to generate diversity within the region linking the two substructures. A final hop is then made in which substructure B in the pair is incorporated into the compound (see the Supplementary Information for example Cypher queries). A schematic demonstrating the key steps in the pipeline is shown in Figure 2.

**Figure 2:**
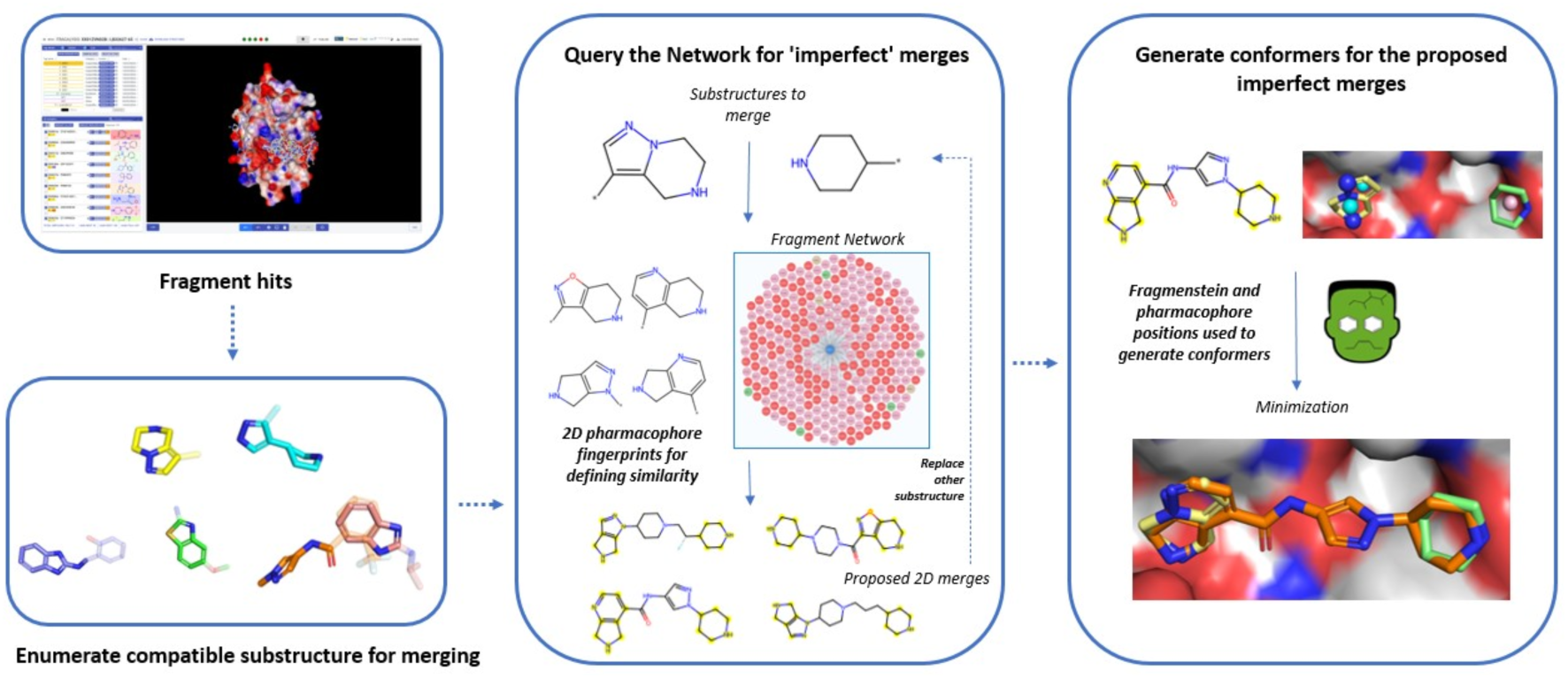
Pipeline for identifying bioisosteric merges. The overall pipeline for using the Fragment Network to find bioisosteric merges begins with the enumeration of compatible substructures from crystallographic fragment hits against a target. Pairs of substructures that are spatially compatible and make interactions with the protein (calculated using ProLIF [30]) are selected. The network is queried to find bioisosteric merges: given a pair of substructures, up to two expansion hops can be made away from the first substructure followed by incorporation of a pharmacophoric equivalent of the second substructure (calculated using similarity between pharmacophoric fingerprints implemented using RDKit [29]). The first substructure can additionally be replaced in subsequent steps. Conformers are generated using an adapted version of Fragmenstein [31]; the replacement substructures are embedded using the pharmacophore coordinates from the original fragment substructures; these are used to generate custom atom-to-atom mappings that are fed into Fragmenstein for embedding the entire structure of the merge against the original fragments. Minimization is performed using PyRosetta [32].

#### 2.4.1 Search for bioisosteric merges

To expand the search to identify ‘bioisosteric merges’, the query follows the same format, except the final expansion step is modified, instead specifying that the substructure to be incorporated need not be identical to the query substructure but should be similar in terms of its pharmacophore features. As described above, the edges in the network are labelled with the pharmacophore fingerprint representing the substructure involved in the transformation. Thus, for a given query substructure, we can calculate similarity to substructures in the database by calculating similarity between their fingerprints, only accepting those with similarity above a specified threshold. The neo4j platform allows this calculation to be performed on-the-fly, enabling automatic filtering of the query results. For the purposes of our search, we used the Tanimoto metric to calculate similarity and retrieved all merges with similarity >0.9 (these query merges can be further prioritised at a later stage, but this threshold was found to result in sufficient numbers of compounds for filtering). This threshold can be adjusted according the requirements of the user.

To further increase diversity within the set of bioisosteric merges, promising compounds can be subjected to a further round of querying following initial conformer generation (Figure 3). To do this, merges for which an alignment has been successfully generated are selected (described below) and used to generate further bioisosteric merges by replacing the seed substructure in the query (also using pharmacophore fingerprint similarity). We provide options for a ‘strict’ and a ‘loose’ search. In the strict search, the seed substructure is removed, no modifications are made to the linker region in the merge and the seed substructure is replaced. In the loose search, an additional contraction and expansion are made within the linker region to generate further diversity and widen the search radius for merges.

**Figure 3:**
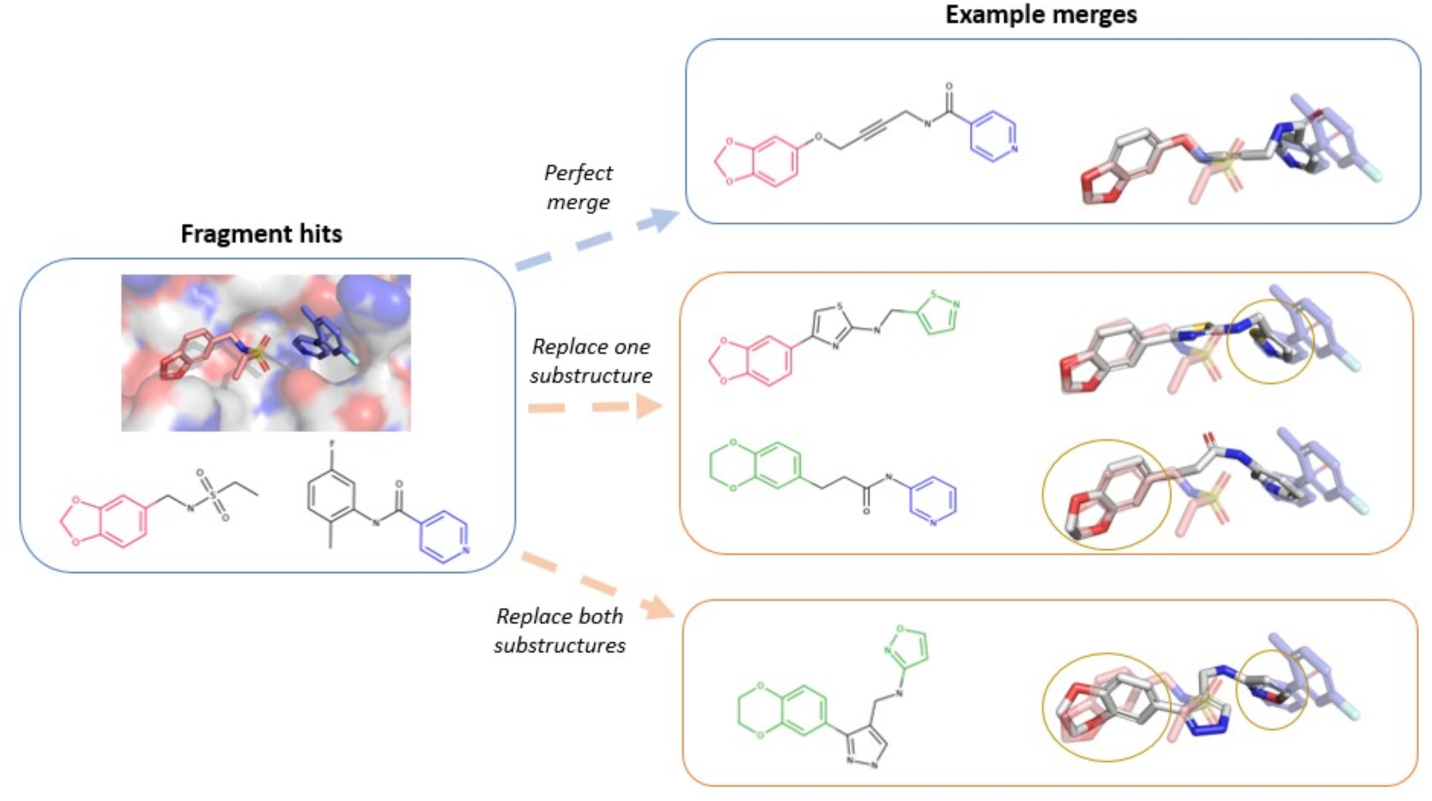
Increasing diversity within fragment follow-up compounds. Diversity within the follow-up compounds can be increased by replacing substructures with those that are ‘pharmacophorically similar’, determined by calculating similarity between pharmacophore fingerprints. Perfect merges maintain the exact substructures (shown in red and blue) seen within the parent fragments. Bioisosteric merges occur when we replace either one or both substructures from the parent fragments (shown in green), increasing chemical diversity within follow-up compounds.

In the results presented here, the results of the perfect and bioisosteric merging pipelines are kept separate to enable comparison (meaning no perfect merges are included in the results of the bioisosteric pipeline). However, when running the tool, the user is able to specify that the search can be expanded to include bioisosteric merges in addition to perfect merges.

As in [5], we also provide the option in the new pipeline to perform R-group expansion at the query stage on the merges to recapitulate substituents seen in the original fragments (this is done in an automated manner requiring no manual curation of possible R-groups). In the results presented, R-group expansion has been additionally run for successfully placed merges.

### 2.5 Prioritization of merges for filtering

Owing to the volume of potential merges retrieved from the database, we prioritised which compounds entered the computationally expensive step of conformer generation. For bioisosteric merges, we extracted all the replacement substructures that are used within the set of merges and generated embeddings using pharmacophores. Pharmacophores are extracted from both the original and replacement substructures and the replacement substructure is aligned to the original pharmacophores according to its 3D coordinates (a process that is relatively computationally cheap). This process results in several possible embeddings (based on which and how many pharmacophores are matched); we used the embeddings that demonstrate the most favourable shape and colour score (using an RDKit implementation, based on the SCr_DKIT_ score described in [33]). The score aims to determine whether binding mode is conserved for elaborated compounds by calculating the overlap between the volume and pharmacophore features between the initial fragments and the elaborated compounds. Following this, possible interactions between the embedded replacement substructure and the protein were calculated (using ProLIF [30]) and compared against the interactions observed for the original substructure. We only maintained merges that incorporate substructures that are observed to make an interaction and ranked them according to the number of potential interactions that are maintained. According to the target and the size of the dataset, we can impose limits on the number of compounds that enter the conformer generation steps per fragment pair. In this work, we imposed a limit of 500 compounds per substructure pair.

### 2.6 Filtering generated merges

#### 2.6.1 Conformer generation

The most important step of our filtering process is to evaluate whether the merges can adopt the conformations observed for the parent fragments. In [5], we used Fragmenstein [31] to generate conformations, which predominantly relies on maximum common substructure matching and positional overlapping between the merge and fragments. Fragmenstein places the merge and can perform minimization within the protein with PyRosetta [32]. As the bioisosteric merges may no longer share exact substructures and may differ in shape (depending on the similarity threshold used), we have adapted the Fragmenstein protocol to perform placement of molecules using pharmacophores rather than molecules. In particular, a fast pharmacophore-based alignment is performed to identify the atomic mapping between the atoms of the fragment and the atoms of the molecule to be placed. This mapping is performed based on distances between atoms, mapping those that are within 1A. The mappings can be provided to Fragmenstein, which subsequently attempts to generate conformations using these mappings; mappings for which Fragmenstein is unsuccessfully able to generate a conformation are discarded. We prioritised the mapped conformers (without minimization) with the highest SCr_DKIT_ score to undergo subsequent minimization within the protein and selected the top 500 compounds to minimise per fragment pair).

Following conformer generation, we apply a series of filters to retain the most promising compounds. We only maintain merges that exhibit a negative ΔΔG value, a combined RMSD of *<2Å* with the substructures used for placement, share overlapping volume with the substructures used for placement and are energetically favourable (according to the energy ratio between several unconstrained conformers and the constrained conformation generated by Fragmenstein, as calculated in the original pipeline [5]).

### 2.7 Comparison against pharmacophore-constrained docking

To compare with more ‘out-of-the-box’ standard techniques for computational follow-up of fragment screens, we also performed docking-based experiments for select fragment pairs. Owing to computational constraints, we did not perform docking of the entire equivalent set of >200 million compounds stored within the database (estimated to require >150,000 CPU days using our docking protocol). We instead extracted representative subsets to minimise the computational costs. Nine fragment pairs were used for comparison, consisting of three fragment pairs across each target that were well-represented amongst the results of the Fragment Network merging pipelines to provide sufficient numbers to enable comparison.

To extract a subset of compounds for docking, we performed a pharmacophore fingerprintbased search with the Tversky metric against every compound in the database, calculating the geometric mean of the similarity against each fragment in the pair. We used fingerprints calculated in a similar way as described above used in annotation of the network substructures using the same pharmacophore features (with the exception of the attachment point atoms). We then extracted the top 100,000 compounds for each pair. To provide a comparison search method that still considers the pharmacophores within the parent fragments, we used rDock [34] to perform a virtual screen of these compound subsets, as rDock provides the ability to dock compounds using a set of specified mandatory and/or optional pharmacophore constraints.

To generate the set of pharmacophore constraints to use for docking, for a given pair of fragments, the pharmacophores that are involved in interactions (calculated using ProLIF [30]) are extracted. The coordinates of the interacting atoms are recorded; for multi-atom pharmacophores, such as aromatic rings, the centroid of the coordinates is calculated. If a pair of pharmacophores are closer than 2Á, one of the pharmacophores in the pair is removed, to ensure that pharmacophores are not too close to prevent finding sensible molecules. The interacting non-hydrophobic pharmacophores were used to generate a set of mandatory pharmacophore constraints. Optional constraints were also imposed, where up to two of the hydrophobic pharmacophores are required to be fulfilled to successfully generate a conformation. A tolerance radius of 2A was used for the pharmacophore alignment. The full set of pharmacophores used for our test cases and the set of parameters used in the docking are provided in Supplementary Tables 3—5.

### 2.8 Case study

To provide validation for our method, we conducted a retrospective case study using hits that were screened against repeat-containing protein 5 (WDR5)-MYC by Chacon Simon et al. [35]. A merging opportunity was identified by overlaying a fragment-screening hit and a hit identified using HTS, for which a compound (compound 2i) was designed with a *K*_d_ value of 1.0^M. The authors also designed several analogues of the merged compound to optimize affinity by exploring several different R-groups. To evaluate whether our bioisosteric pipeline is able to replicate some of the design ideas used in the optimization and thus could act as a useful tool for early SAR exploration, we ran our merging pipeline and provide comparison of the outputted compounds against the existing designed compounds with experimental data, consisting of *K*_d_ values from a fluorescence polarization-based assay.

## 3 Results

### 3.1 Enumeration of substructure pairs for merging

We tested our bioisosteric and new perfect merging pipelines using crystallographic fragment hits against three different antiviral targets: EV D68 3C protease, EV A71 2A protease and Zika NS2B.

For the EV D68 3C protease, we considered 25 fragment hits found to bind to the catalytic site of the protein. After running our enumeration pipeline (where we removed pairs of fragments according to overlap and remove pairs of substructures according to overlap and whether they are responsible for making an interaction), we found 152 pairs of fragments (querying is asymmetric; thus, the same fragment pair may be represented twice), leading to 1,128 pairs of substructures for querying. Totals for numbers of enumerated pairs are in Table 1. Timings for querying and conformer generation can be found in Supplementary Tables 6,7.

**Table 1:** Enumerated pairs of substructures for merging.

| Target | Number of fragment hits | Enumerated number of fragment pairs* | Enumerated number of substructure pairs |
| --- | --- | --- | --- |
| EV D68 3C protease | 25 | 152 | 1,128 |
| EV A71 2A protease | 6 | 11 | 88 |
| Zika NS2B | 41 | 169 | 1,026 |

*\*Querying is asymmetric; thus, a fragment pair may be repeated. EV, enterovirus.*

In the case of the EV A71 2A protease, at the time we ran the pipeline (December 2023), only the results of a partial fragment screen were available, meaning we focused on only six compounds found to bind to the main active site of the target. This is a valuable test case to include as it reflects the realistic setting of drug development for a target, where there may be several iterations of data release and follow-up compound design. The enumeration pipeline resulted in 11 fragment pairs and 88 pairs of substructures for querying.

For Zika NS2B, due to the dense population of fragments within the S1 site (with 37 fragments bound; Figure 1c), we focused only on merges that occur between subsites (that is, between S1 and S1‘ or between S1 and S2) to find molecules that bridge across multiple sites. This enumeration resulted in 169 fragment pairs leading to 1,026 pairs of substructures for querying.

### 3.2 Bioisosteric merging identifies potential follow-up compounds across all targets

Fragmenstein was used for generating conformations for our 2D proposed merges based on the parent fragments. Substantial redundancy was observed across the compound set as the same compound may be suggested for different pairs of fragments; filtering was run for each instance as the coordinates of the specific fragment may affect the probability of generating a successful alignment. Supplementary Figure 3 shows the numbers of unique filtered compounds that were successfully placed using Fragmenstein and passed our filters for each merging pipeline (all values for how many compounds pass through each step are shown in Supplementary Table 8). Comparable numbers of compounds are identified when comparing the collective results from both rounds of bioisosteric merging with the results of perfect merging (across all targets). The efficiency of the search (defined as the percentage of compounds that enter minimization that are successfully placed with a SC_rDKIT_ value of ≥0.55; see Methods for definition) ranged between 0.7-3.3%, with the lowest efficiency observed for the second round of bioisosteric merging for the EV A71 2A protease.

Upon visual inspection, the bioisosteric merges show a high degree of volume overlap with the fragments used for inspiration and many of the merges are observed to have high shape similarity with the original fragments (particularly where replacement substructures maintain the rings seen in the original substructures), which indicates that bioisosteric merging still shows promise for recapitulating the original fragment poses. Examples of bioisosteric merges are shown in Figure 4, in which the computationally predicted structures of the merges mirror the structures of the crystal fragments and are predicted to maintain interactions seen in both of the parent fragments.

**Figure 4:**
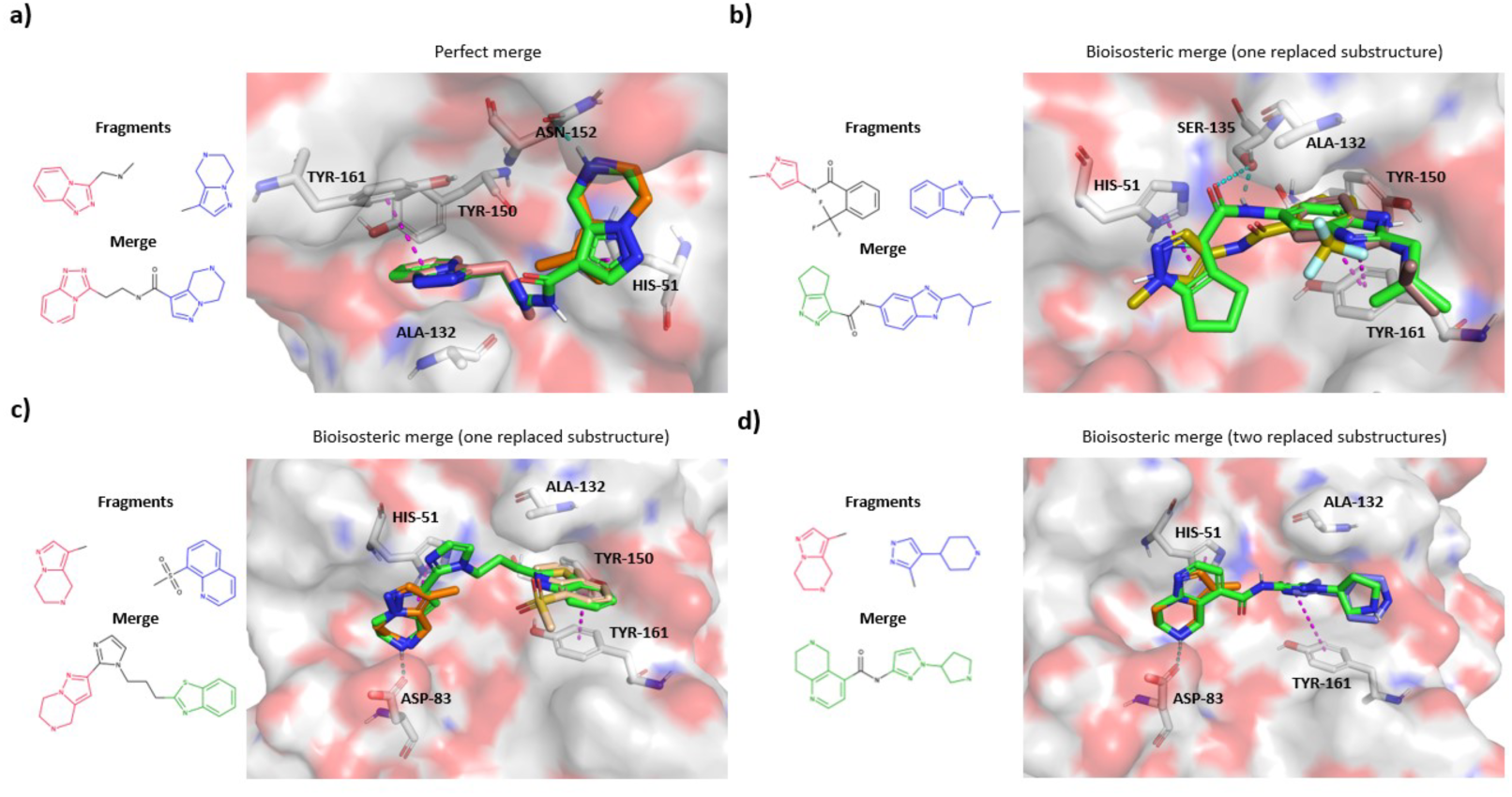
Merges identified for fragment hits against Zika NS2B. Example merges against Zika NS2B are shown (Fragmenstein-predicted structures in green), overlaid with the crystallographic fragment hits used as inspiration for their design. The Fragment Network is used to identify (**a**) perfect merges, which contain exact substructures from two fragment hits, (**b,c**) bioisosteric merges that incorporate one replaced substructure and one exact substructure from the fragments and (**d**) bioisosteric merges that incorporate two replaced substructures. Residues predicted to be involved in interactions are shown. Hydrogen bonds are shown in cyan and n-stacking interactions are shown in magenta; hydrophobic interactions are not shown.

### 3.3 Bioisosteric merging expands the representation of fragment pairs

In the following sections, we compare various metrics across compounds found via bioisosteric merging (after both rounds of bioisosteric merging) versus perfect merging. To rule out compounds that do not represent favourable merges, the analysis is limited to compounds that have a SCr_DKIT_ score ≥0.55 (this threshold indicates a favourable degree of volume and pharmacophore feature overlap [36]) and that are predicted to replicate an interaction observed for each original fragment (thus representing true merges). To remove redundant compounds (with the same SMILES), compounds are ordered according to their SCr_DKIT_ and the SMILES with the highest SCr_DKIT_ scores are maintained. The total numbers of compounds and their observed features are shown in Figure 5. Across all targets, bioisosteric merging is found to represent fragment pairs that were not represented by perfect merging and, for EV D68 3C protease and Zika NS2B, perfect merging found merges for pairs not represented by bioisosteric merging. When comparing the number of individual fragments represented, for EV D68 3C protease, both methods represent features from the same 16 fragments, while, for the EV A71 2A protease, bioisosteric merging identifies merges for 2 fragments not shown in the perfect merging set and both methods for Zika NS2B identify merges for a fragment not represented by the other. Bioisosteric merging is, therefore, valuable for expanding the search space of the Fragment Network search and identifies opportunities for fragments that are not well-represented by merging exact substructures alone.

**Figure 5:**
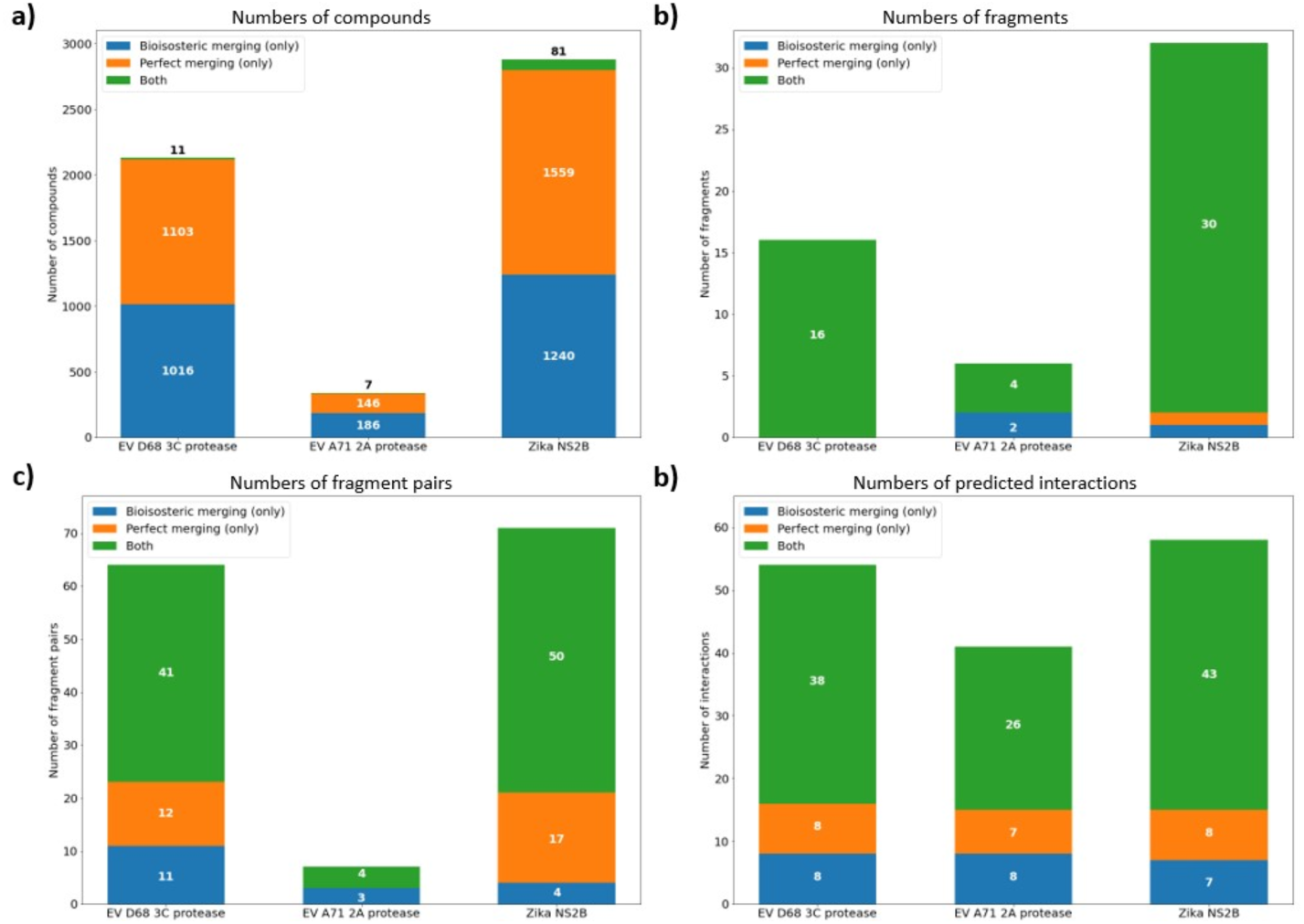
Numbers of fragments, pairs of fragments and predicted interactions represented by merges found using perfect or bioisosteric merging. For the top-performing merges (SCrdkit score >0.55 and are predicted to maintain an interaction seen for both parent fragments), we show (**a**) the number of compounds identified, (**b**) the number of individual fragments used in the design of the compounds, (**c**) the number of fragment pairs represented by the compound sets and (**d**) the total numbers of unique interactions predicted to be made by the entire compound sets. Colours indicate whether the compound, fragment, fragment pair or interaction was identified by compounds only retrieved via perfect merging, bioisosteric merging or both methods.

### 3.4 Bioisosteric merging identifies potential interactions not represented by perfect merging

When we compare the unique interaction types that are predicted to be made by merges proposed by the bioisosteric versus perfect merging pipelines referring to the number of residue-level interactions made by each compound within the set), the results suggest that bioisosteric merging does not degrade the ability of the method to find merges that replicate the observed interactions. Bioisosteric merges perform similarly in terms of the mean total number of interactions made, dependent on target, being marginally higher (as observed for EV D68 2A protease and EV A71 2A protease) or slightly lower (for Zika NS2B) (Supplementary Figure 4). Similarly, when considering the proportion of preserved interactions, no technique performs consistently better than the other (Supplementary Figure 5).

Figure 5d shows, for each target, the number of interactions predicted to be made across the compound sets that are only observed for either the perfect or bioisosteric pipelines. Depending on the target, bioisosteric merges are found to identify between 7 and 8 interactions that are not represented with perfect merging alone, demonstrating that this method is useful for expanding the potential interaction space explored.

### 3.5 Bioisosteric merging represents more unique substructures than perfect merging

Supplementary Figure 7 shows the number of unique substructures that are found among merges from both search techniques (that is, the number of substructures either taken directly from the fragments or that have been proposed as bioisosteric replacements, defined according to the SMILES string). Bioisosteric merging results in a substantial increase in the number of substructures represented, showing a ∼2.4-5.3-fold increase. This demonstrates how bioisosteric merging can expand the search space and thus, potentially, the chemical diversity seen in the final compounds (see Supplementary Figure 8 for examples).

### 3.6 Bioisosteric merging increases the chemical space coverage of the search

One aim of bioisosteric merging is to access new areas of chemical space that are not explored when considering only perfect merges. Figure 6 shows low-dimensional representations of chemical space using T-SNE plots [37]. For all targets, the plots show a separation between the regions of chemical space that are occupied by bioisosteric and perfect merges. There are clusters of compounds for bioisosteric merges that are not accessed by the perfect merge search (as well as some overlapping regions). To explore this further, we clustered the outputs of bioisosteric and perfect merging for each target (Supplementary Table 9), which shows the results of clustering using the Butina algorithm [38] with two different distance thresholds (calculated using Tanimoto similarity between Morgan fingerprints). Overall, this analysis showed that far fewer clusters contain compounds proposed by both techniques and bioisosteric merging was able to identify hundreds of clusters that contain no perfect merges, indicating that the method expands the chemical space searched.

**Figure 6:**
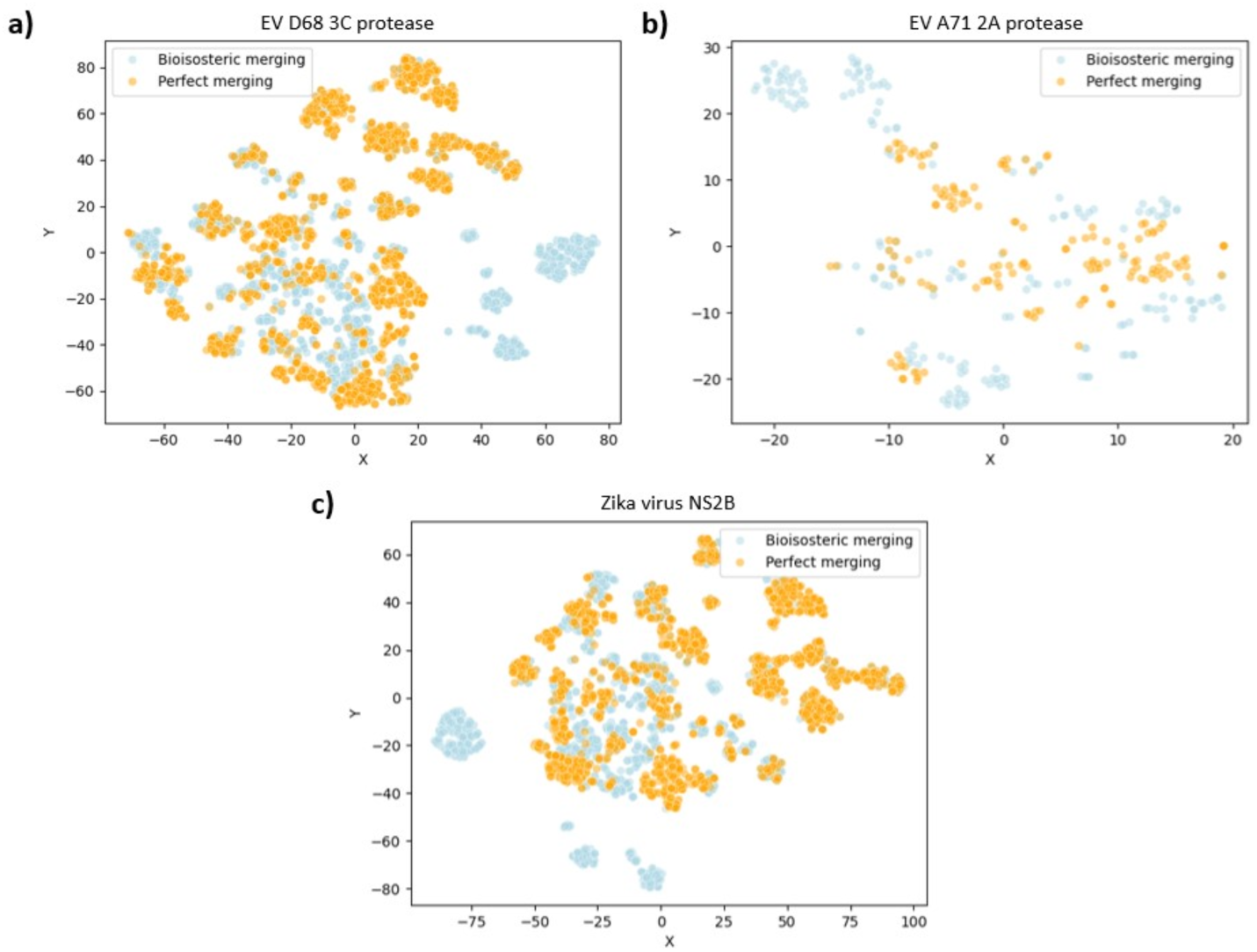
Chemical space coverage by Fragment Network merges. T-SNE plots depict lowdimensional representations of chemical space for filtered merge compounds identified for targets (**a**) enterovirus (EV) D68 3C protease, (**b**) EV A71 2A protease and (**c**) Zika virus NS2B. Molecules are represented using Morgan fingerprints of length 1,024 bits and a radius of 2. The dimensionality of the vectors was reduced to 50 using PCA before applying the T-SNE.

### 3.7 The Fragment Network merging pipeline offers potential efficiency benefits over pharmacophore-constrained docking

To enable comparison with an out-of-the-box docking approach, we performed pharmacophore-constrained docking using rDock for nine pairs of fragments, consisting of three fragment pairs against each protein target. The pharmacophore constraints were selected according to pharmacophore features that were responsible for making interactions (calculated using ProLIF; constraints are provided in Supplementary Tables 3—5). For each merge that passed through the rDock pipeline, the conformer with the best docking score was selected. The SCrdkit scores are consistently higher across all pairs for Fragment Network-identified compounds (Supplementary Figure 9), which can be attributed to several factors: the Fragment Network approach aims to maintain atoms (or similar substructures) directly inspired by the original fragments, but also the conformation generation method (using Fragmenstein) deliberately attempts to align the compounds with the parent fragments as closely as possible, whereas rDock conformers are ranked according to its scoring function (example conformations for both methods are shown in Figure 7). Thus, to enable comparison between methods, we employed a less conservative SCr_DKIT_ threshold by filtering out compounds with a SCr_DKIT_ score of less than 0.4. Compounds are instead ranked according to the frac-tion of preserved interactions (meaning the proportion of unique residue-level interactions made by the parent fragments maintained by the merge); comparisons in Supplementary Figures 10-16 and Supplementary Table 10 are made between the top 100 compounds from both methods.

**Figure 7:**
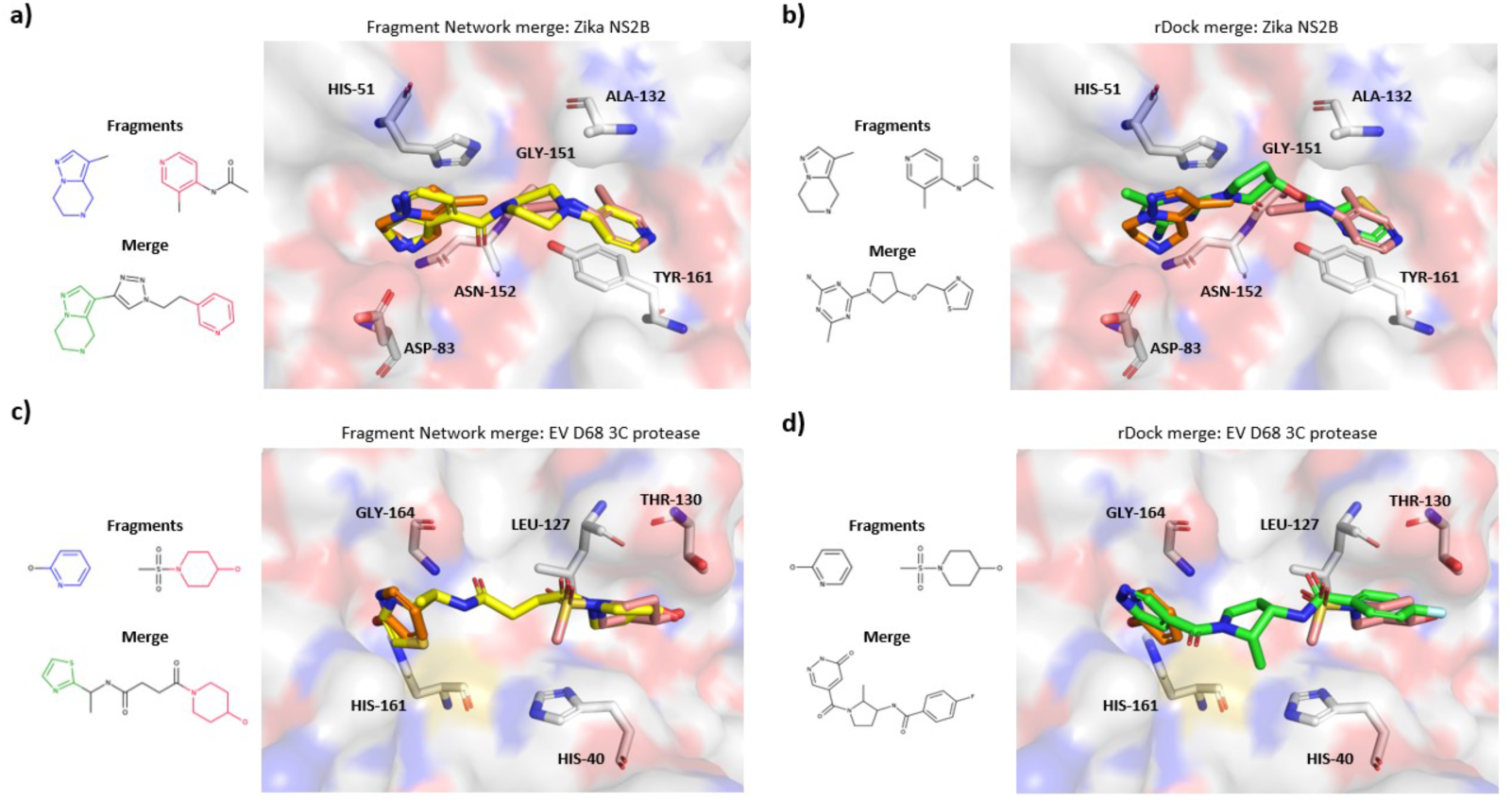
Examples of high-scoring merges from bioisosteric merging and rDock pipelines. We show example high-scoring merges (according to shape and colour scores) for bioisosteric merging (using the Fragment Network) pipelines and using pharmacophore-constrained docking using rDock. Examples against a fragment pair against Zika NS2B (**a,b**) and enterovirus (EV) D68 3C protease (**c,d**) are shown. Crystallographic fragment hits are shown in orange and pink. Computationally predicted conformers of Fragment Network compounds are shown in yellow and rDock-identified compounds are shown in green.

**Figure 8:**
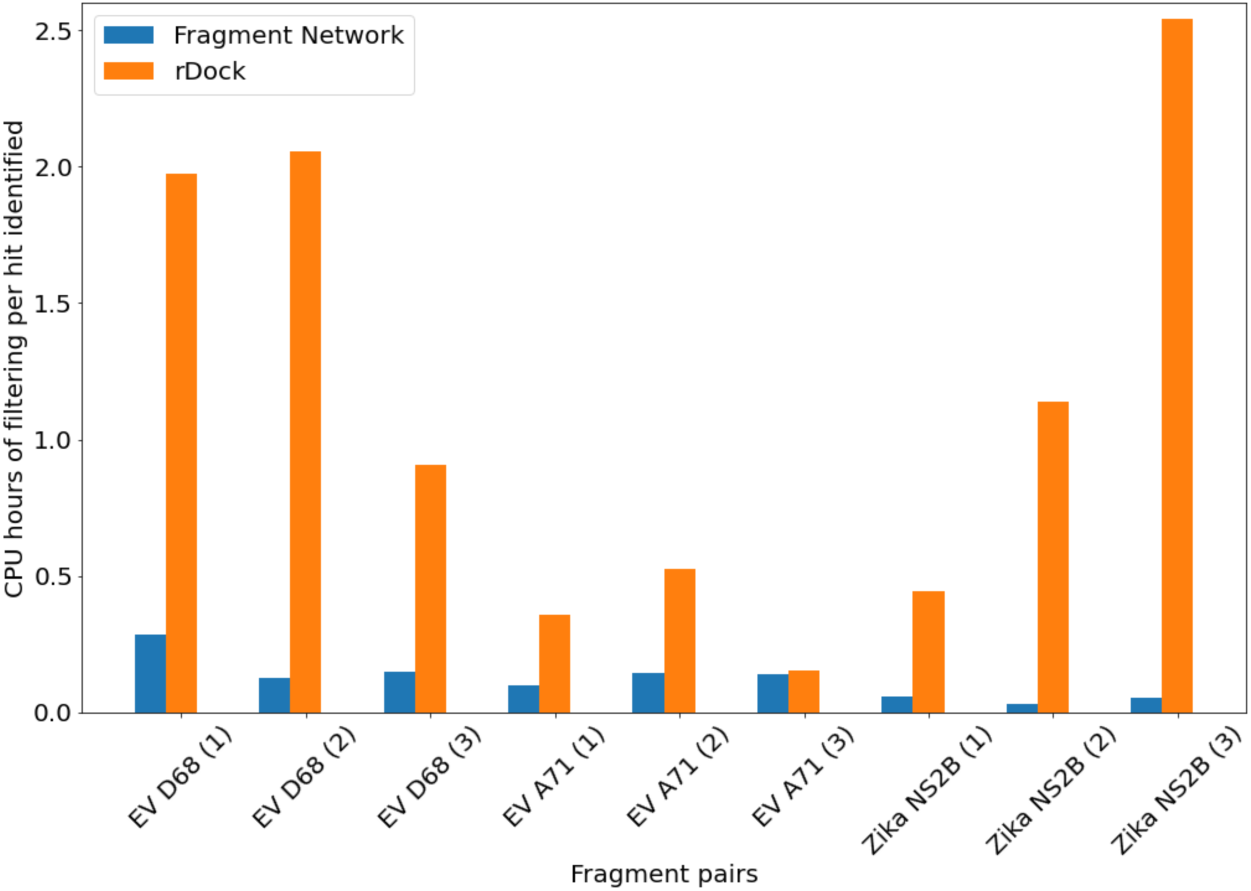
CPU hours of filtering required per computational merging ‘hit’ identified for pharmacophore-constrained docking versus Fragment Network merging. The CPU hours required for conformer generation to identify a high-scoring merging ‘hit’ are shown using our Fragment Network merging pipelines (using 3D placement and minimisation with Fragmenstein) versus pharmacophore-constrained docking following a similarity search using a pharmacophore fingerprint. The ‘hits’ are defined as compounds that preserve >50% interactions observed in the parent fragments and demonstrate a mean SCr_DKIT_ of at least 0.4.

When comparing the chemical space coverage of the two techniques, it is noticeable across all pairs that the molecules identified by each technique occupy distinct areas of chemical space. Clustering analysis also shows that the docking-identified compounds are more chemically diverse, occupying more clusters across all pairs of fragments than Fragment Network-identified compounds, and no clusters were found to include compounds from both techniques (Supplementary Table 10). Regarding the interactions that are predicted to be made by the follow-up compounds, neither technique identifies compounds that are predicted to make a greater number of interactions consistently across all pairs (including when normalised according to heavy atom count; Supplementary Figures 10 & 11).

To compare the efficiency of the two search approaches, we evaluated the ability of the two methods to identify computational potential merging ‘hits’; we have defined this as compounds with a SCrdkit score of at least 0.4 and that preserve at least 50% of unique residue-level interactions observed for the fragments. We chose this SCr_DKIT_ score following visual inspection of the merges, as we found this score was still sufficient enough for there to be a plausible hypothesis as to which structures from the fragments were used as inspiration, whereas this was not always evident with lower scores for the docked compounds. While true evaluation of whether compounds constitute a hit requires experimental validation in binding assays, this metric provides a general indication of the resources required to identify the most promising compounds that could contribute to a potential shortlist for purchase or synthesis. Table 2 shows, per fragment pair, the number of compounds (and associated CPU hours) required to yield the number of identified computational merging hits. While the docking protocol performs favourably for certain pairs, in particular for those against the EV A71 2A protease 8), the Fragment Network pipeline requires less resources in terms of the CPU hours required per hit across all pairs (between a 1.1-49.5-fold reduction in hours). We hypothesise that the improved performance of the docking protocol for EV A71 2A protease may be due to the increase in overlapping volume between the fragments, meaning there are fewer (and closer) pharmacophores provided as docking constraints.

**Table 2:**
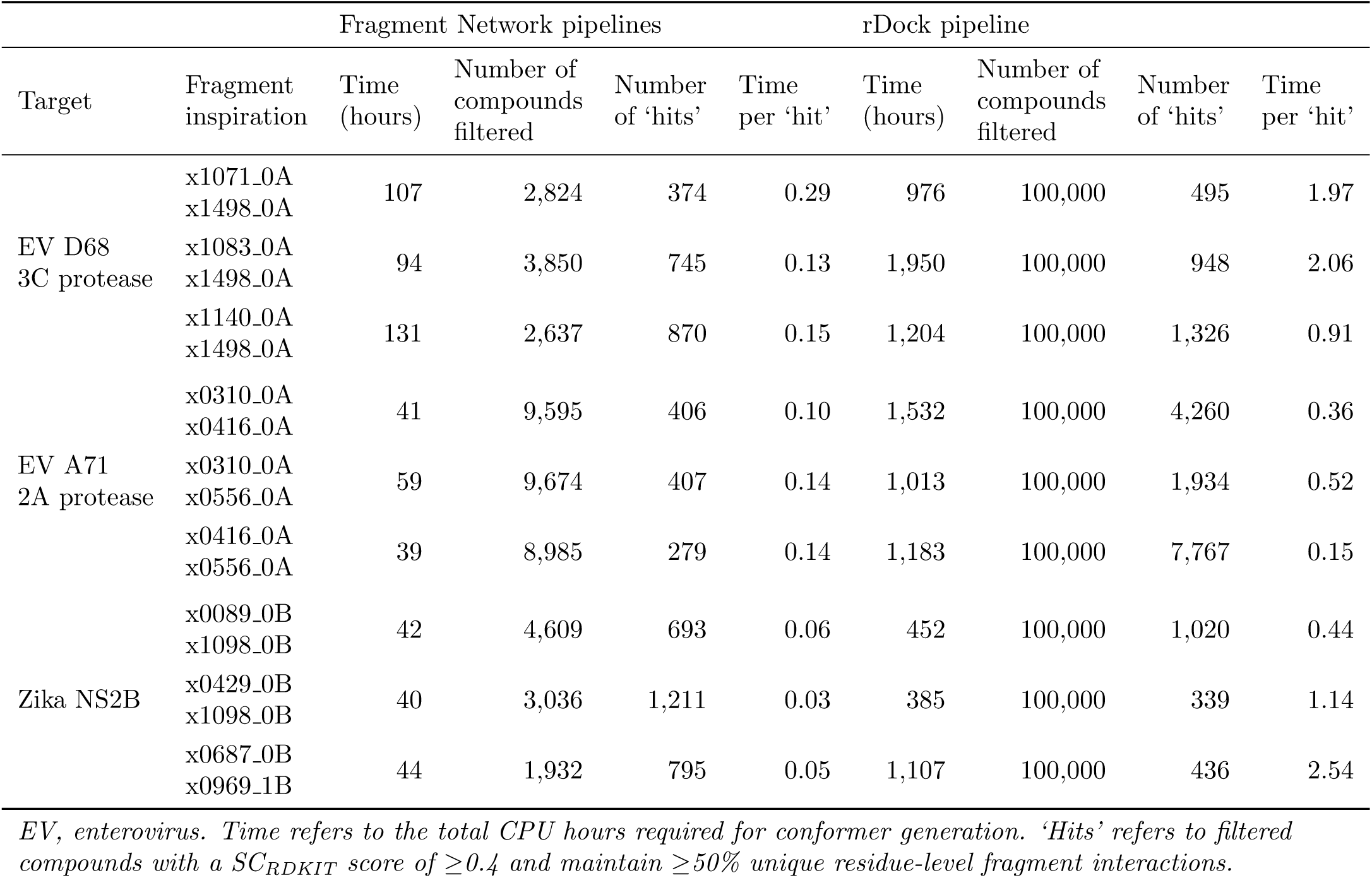
Pharmacophore-constrained docking versus Fragment Network merging.

These results indicate that — while constrained docking can be a powerful method for follow-up development in certain contexts and does not have the same memory requirements for implementing the Fragment Network infrastructure — either larger numbers of molecules have to be screened, necessitating substantial compute, or more stringent pre-filtering methods are needed to reduce the library size to manageable levels. While the docking approach is adept at identifying highly diverse follow-up designs, the docking experiments conducted were not sufficient to consistently identify more compounds that maximise interactions across all of the targets when scaled by computational time. While both pipelines could be further optimised, and experimental validation is needed to fully assess the performance of each method, the ability of our tool to identify a comparable number of promising compounds with substantially less filtering time indicates it is well-suited to scenarios in which we want to quickly leverage all potential merging opportunities from a fragment screen.

### 3.8 Case study: our pipeline replicates design choices made during SAR analysis

To provide retrospective validation using existing experimental results for manually designed fragment merges, we compared designs proposed by our perfect and bioisosteric merging pipelines with the results of inhibitors designed against WDR5-MYC [35]. In the original paper, the authors designed and optimized a merge of a fragment-screening hit (referred to as F2), and a hit identified by HTS (compound 1). The merge incorporates features from both hits and performs some additional optimization around the benzene ring of compound 1 through the addition of a hydroxyl group and a chlorine atom (Figure 9a,b). Several analogues were also designed that build a picture of the SAR by exploring different R-groups around the molecule.

**Figure 9:**
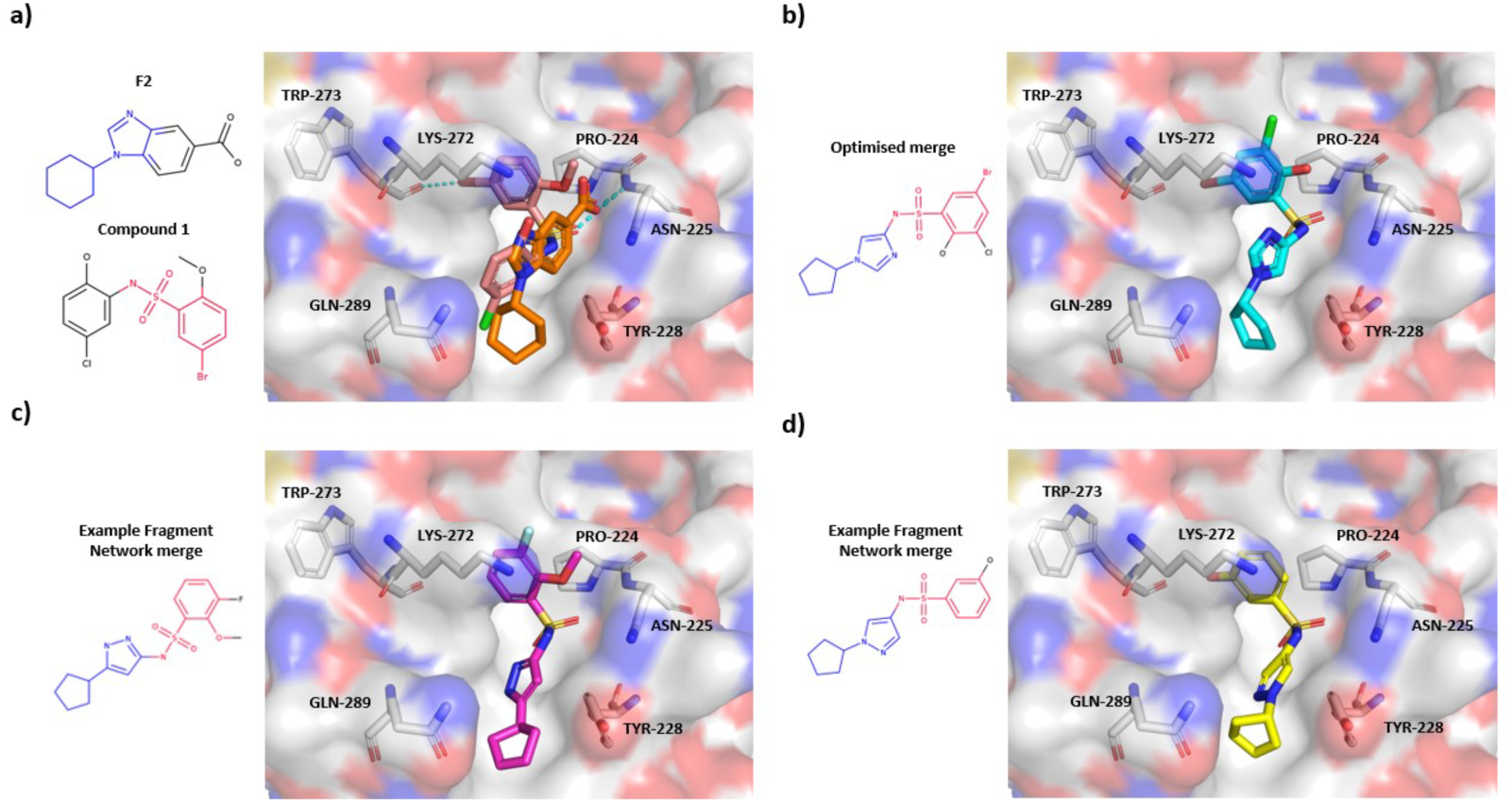
Case study involving merges designed against WDR5-MYC. (**a**) Crystal structures of two hits identified by fragment screening (compound F2; orange) and high-through screening (compound 1; pink) against WD repeat-containing protein 5 (WDR5)-MYC that were used as inspiration for merge design [35]. (**b**) Crystal structure of the original manually designed merge; further optimization is performed through substituting in a hydroxyl group and addition of a chlorine atom. (**c,d**) Example merges proposed by the Fragment Network pipeline that replace the imidazole ring seen in the original fragment with pyrazole rings with different arrangements of nitrogen atoms. Conformations have been generated using Fragmenstein [31].

In the original work [35], the analogue design process included the addition of alternative heterocycles to the imidazole ring from hit F2; there were two transformations whereby a pyrazole ring was added with different arrangements of nitrogens around the ring (referred to as substituents 3b and 3d).

We ran our pipeline to find catalogue merges incorporating substructures from F2 and compound 1. The initial parameters for filtering substructure pairs were loosened, such as removing substructures composed of single carbon rings from querying, due to their presence in the parent compounds and as we do not face the same computational limitations when only searching for merges for a single pair of fragments (these parameters can be freely modified by the user according to their requirements). We find that our pipelines proposed several merges that incorporate these pyrazole rings while maintaining the cyclopentane ring (found to have favourable effects for binding affinity in the original paper [35]). This included 12 compounds that incorporate the pyrazole ring arranged as in substituent 3b, and 6 compounds that incorporate the pyrazole ring arranged as in substituent 3d. Our pipelines also identify compounds that incorporate various substituents added to the benzene and pyrazole rings, which may provide further opportunities for elucidating SAR and suggesting potential optimizations. We include an R-group decomposition representing these proposed molecules and the various added substituents in Supplementary Figures 17 & 18.

In addition to being able to recapitulate some of the manual design ideas, Figures 9c,d, show how the generated conformers demonstrate reasonable alignment with the crystallographic structures of the original fragments and the manually designed merge. This demonstrates how some of the substitutions incorporated in our bioisosteric merging pipeline can begin to mirror some of the decisions made by chemists when exploring SAR for a given target.

## 4 Discussion

In this work, we describe an update to our method for merging fragments using the Fragment Network that maximises chemical diversity by incorporating substructures that recapitulate pharmacophoric features from the original fragments. Our bioisosteric merging method widens the search space for follow-up compounds, potentially improving the productivity of the catalogue search. Results are shown across multiple viral targets representing vastly different merging opportunities with regard to the numbers of fragment hits observed to bind, the size of the pockets and the distribution of fragment hits in pocket space, demonstrating the robustness of the pipeline and its suitability in multiple scenarios. Our tool aims to provide a more efficient search approach than traditional docking-based methods, requiring less computational resources and thus making it suitable for situations where we want to exploit all possible elaboration opportunities from a fragment screen.

When selecting compounds for purchase, a major consideration is maintaining as much diversity as possible and representing multiple core scaffolds in order to spread risk and increase the likelihood of finding a chemical series that can bind. Thus, it is promising that the bioisosteric merging pipeline is able to identify clusters of compounds in distinct areas of chemical space, representing up to a 5.3-fold increase in the number of unique substructures incorporated.

Whilst delivering this increased diversity, the bioisosteric merging pipeline showed comparable performance with our original perfect merging in terms of the numbers of interactions predicted for a compound, the number of new unique interactions found that were not seen in the original fragments and the fraction of preserved interactions. Increased diversity is also reflected in terms of the observed interactions, as, across all targets, bioisosteric merges were able to identify potential new unique interaction types that were not observed using perfect merging alone. We also find that the generated conformers show favourable pharmacophoric overlap with the parent fragments, although many of the merges already share a high degree of shape similarity owing to the replacement substructures used.

We compared our new approach with a more standard out-of-the box approach based on pharmacophoric similarity search and constrained docking, which resulted in small differences in terms of the numbers of total potential interactions that can be exploited. To provide a method of comparison, we offer a definition of a computational merging ‘hit’, referring to compounds that demonstrate a SCr_DKIT_ score of at least 0.4 and maintain at least 50% of the interactions observed within the parent fragments. The classical approach did sample a more diverse set of compounds but required substantially more compute when comparing the hours of conformer generation needed per merging ‘hit’ identified. These results suggest our method may be more convenient when scaling up to searching for merges and linkers of all possible fragment pairs. While the definition of a merging hit can be subjective and requires further experimental validation, our pipeline enables efficient identification of these favourably scoring compounds.

Overall, our tool provides increased chemical diversity compared with the previous iteration and provides a step towards the automation of SAR analysis for merges in a much more efficient way than previous techniques.

## Supporting information

Contains all supplementary material referenced in the manuscript.

## 5 Code availability

The code for running the Fragment Network merging pipelines is available from https://github.com/stephwills/FragmentKnitwork. Data for this work are available from https://doi.org/10.5281/zenodo.13151113. Data files for running this version of the Fragment Network database will be made available.

## Acknowledgements

The authors would like to thank T. Dudgeon and A. Christie for their help with setting up the Fragment Network database. The authors would also like to thank M. Ferla for help and advice with running Fragmenstein. This work was supported by the Engineering and Physical Sciences Research Council (EPSRC; grant number EP/S024093/1), Vernalis (R&D) Limited, and LifeArc.

