## Supplementary material for "Expanding the scope of a catalogue search to bioisosteric fragment merges using a graph database approach": Contains all supplementary material referenced in the manuscript.

### 7 Supplementary Information

#### 7.1 Crystallographic data

Table 1: Crystallographic fragment screening hits used for merging.

| Target | Fragments |
| --- | --- |
| Enterovirus D68 3C protease | **P1**: x0147_0A, x0771_0A, x0789_0A, x0980_0B, x1332_0A, x1498_0A, x1537_0A, x1594_0A, x1604_0A, x2099_0A, x2135_0A, x2148_0A, x2149_0A, x1163_0A<br>**P2**: x0130_0A, x1020_0A, x1083_0A, x1084_0A, x1140_0A, x1247_0A<br>**P2 & P3**: x1071_0A, x1329_0A, x1919_0A, x2021_0A, x1052_0A |
| Enterovirus A71 2A protease | x0451_0A x0554_0A x0556_0A x0566_0A x0310_0A x0416_0A |
| Zika NS2B | **S1**: x0051_0B x0089_0B x0101_0B x0182_0B x0227_0B x0229_0B x0382_0B x0386_0B x0422_0B x0425_0B x0429_0B x0435_0B x0443_0B x0455_0B x0465_0B x0472_0B x0490_0B x0553_0B x0559_0B x0589_0B x0605_0B x0680_0B x0687_0B x0693_0B x0727_0B x0773_0B x0788_0B x0800_0B x0803_0B x0852_0B x0884_0B x0904_0B x0917_0B x0945_0B x0951_0B x0969_0B x0990_0B<br>**S1′**: x0846_0B<br>**S2**: x0404_0B x0969_1B x1098_0B |

#### 7.2 Updates to the Fragment Network

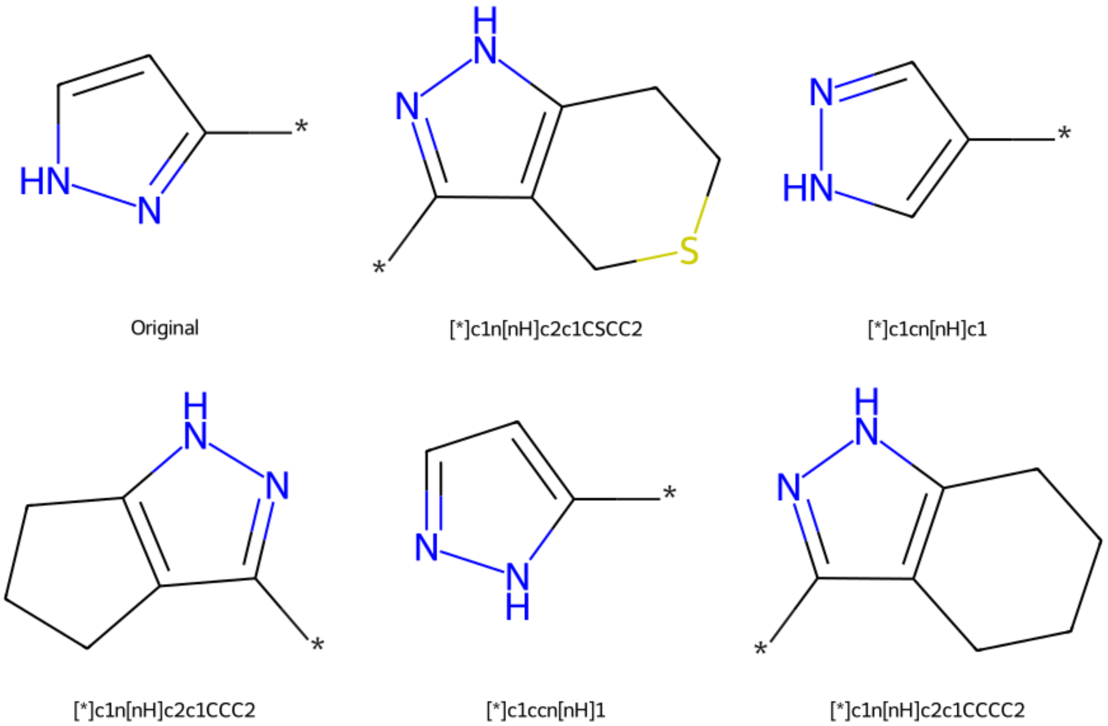


Figure 1: **Example substructure replacements used in bioisosteric merging.** An example substructure from a crystallographic fragment hit (top left) and five example replacement substructures found in the Fragment Network. The asterisk denotes the attachment point atom. Some replacement substructures may be chemically identical to the original but differ in their attachment point.

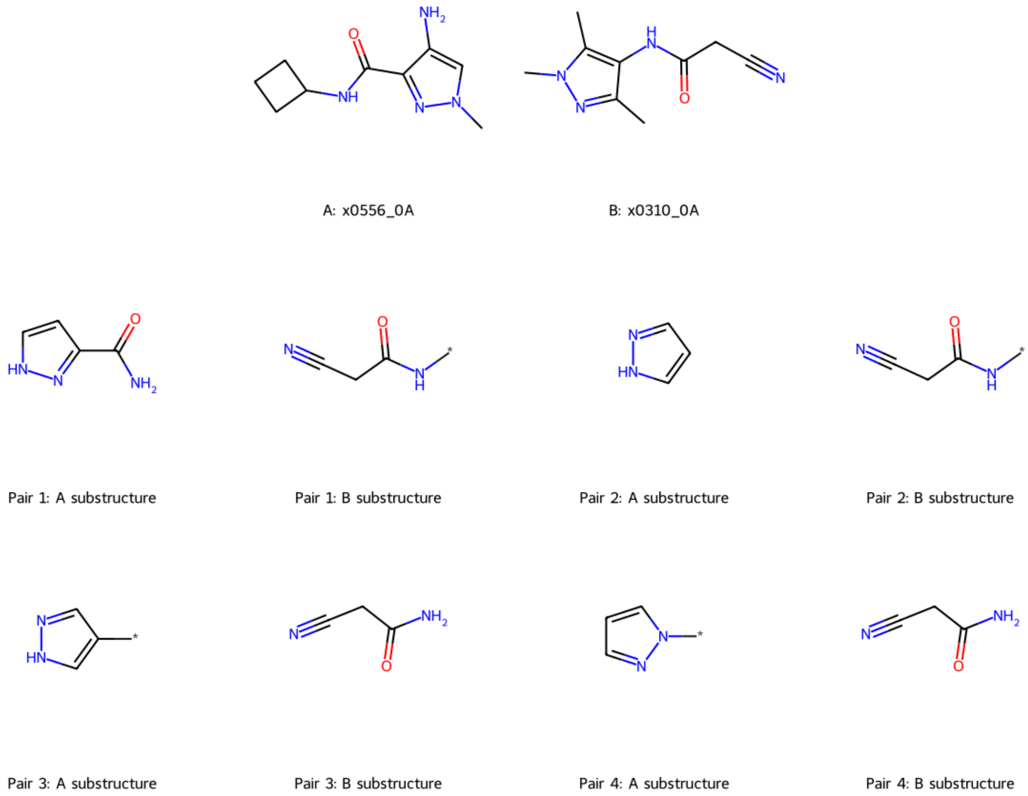


Figure 2: **Example pairs of compatible substructures for merging.** Enumerated substructure pairs for querying are shown for fragment hits x0556-0A and x0310-0A against enterovirus A71 protease 2A. The substructures have been filtered according to their degree of overlap (<30% the volume of one fragment) and as to whether they make an interaction (calculated using ProLIF).

#### 7.3 Example Cypher queries

Example Cypher queries are shown below for the different types of merging. The queries have been simplified for illustrative purposes. Values within inequality symbols (<VALUE>) indicate user defined parameters and values. Colons are used to represent labels (of specific node or edge types): :Mol represents nodes that are purchasable molecules; :F2 represent all molecule nodes (i.e., whether they represent purchasable molecules or intermediate transformations between purchasable molecules), and :FRAG represents edges whereby a transformation (a contraction or expansion) is being made.

**Perfect merging**

Below is an example Cypher query for finding a perfect merge. First, the node representing the seed substructure is identified (<SMILES>). Following this, up to two hops are made whereby expansions occur, which generates diversity ([:FRAG*0..2]), followed by an additional hop ([e:FRAG]), to identify a purchasable catalogue compound (c:Mol). The expansion specifies that the query substructure (<QUERY_SUBSTRUCTURE>, representing the other substructure in the pair) is to be incorporated, based on SMILES matching. The SMILES of the purchasable molecule (representing the merge) is returned on the final line (RETURN molecule_smiles).

```
MATCH (a:F2 {smiles: <SMILES>})<-[:FRAG*0..2]-(b:F2)<-[e:FRAG]-(c:Mol)
WHERE e.substructure = <QUERY_SUBSTRUCTURE>
WITH c.smiles as molecule_smiles
RETURN molecule_smiles
```

**Bioisosteric merging: round 1**

Below shows an example Cypher query for finding a bioisosteric merge. The node representing the seed substructure is identified (<SMILES>). As above, up to two hops are made whereby expansions occur ([:FRAG*0..2]), followed by an additional hop ([e:FRAG]) to identify a purchasable catalogue compound (c:Mol). Similarity is calculated between the pharmacophore fingerprint of the substructure incorporated in the final hop (e.pharmfp) and a query pharmacophore fingerprint, representing the other substructure in the pair (<QUERY_PHARMFP>), while ensuring merges are not retrieved with the exact original substructure (which is achieved by perfect merging). The SMILES of the final purchasable molecule is returned on the last line (RETURN molecule_smiles).

```
MATCH (a:F2 {smiles: <SMILES>})<-[:FRAG*0..2]-(b:F2)<-[e:FRAG]-(c:Mol)
WHERE EXISTS(e.pharmfp)
WITH similarity.tanimoto_similarity(e.pharmfp, <QUERY_PHARMFP>) as tanimoto,
c.smiles as molecule_smiles
WHERE tanimoto >= <THRESHOLD>
AND NOT e.substructure = <QUERY_SUBSTRUCTURE>
RETURN molecule_smiles
```

**Bioisosteric merging: round 2**

Below shows an example Cypher query for performing an additional round of bioisosteric merging. The SMILES of the first merge is matched (<SMILES>), which then undergoes a contraction to remove the original seed substructure in the query ([e1:FRAG]) followed by an additional contraction ([e2:FRAG]) and expansion ([e3:FRAG]) to generate diversity in the linker region linking the two substructure. (This step can be skipped to allow a stricter search.) Following this, an expansion is made in which a similar substructure is incorporated ([e4:FRAG]). The WHERE clauses set constraints on the query, ensuring that the same substructure isn't removed and re-added, and that the substructure added in the first round of bioisosteric merging isn't removed.

```
MATCH (a:F2 {smiles: <SMILES>})-[e1:FRAG]->(b:F2)-[e2:FRAG]->(c:F2)
<-[e3:FRAG]-(d:F2)<-[e4:FRAG]-(f:Mol)
WHERE e1.substructure = <QUERY_SUBSTRUCTURE>
AND EXISTS(e4.pharmfp)
AND NOT e2.substructure = <FIRST_SUBSTRUCTURE>
AND NOT e2.substructure = e3.substructure
AND NOT e3.substructure = e1.substructure
WITH similarity.tanimoto_similarity(e1.pharmfp, e4.pharmfp) as tanimoto,
f.smiles as molecule_smiles
WHERE tanimoto >= <THRESHOLD>
RETURN molecule_smiles
```

##### 7.4 Features used for pharmacophore-constrained docking

Docking parameters used for docking with rDock are shown in Table 2. Mandatory and optional pharmacophore constraints used for select fragment pairs are shown in Tables 3–5.

Table 2: Parameters used for docking with rDock

| Parameter | Value |
|---|---|
| Type of cavity method | Reference ligand |
| Cavity radius | 8.0 |
| Number of docks | 30 |
| Pharmacophore weight | 1 |
| Tolerance radius | 2.0 |

Table 3: Pharmacophore features for docking merges against EV D68 3C protease

| Pair | Requirement | x | y | z | Type |
|---|---|---|---|---|---|
| x1071_0A-x1498_0A | Mandatory | -7.09 | -4.283 | -3.9 | Acc |
| | Mandatory | -7.115 | -1.971 | -2.213 | Don |
| | Mandatory | -5.876 | -1.261 | 0.271 | Acc |
| | Mandatory | -6.835 | -3.99 | -9.255 | Acc |
| | Optional | -7.907 | 0.111 | -0.552 | Hyd |
| | Optional | -4.912 | -7.449 | -2.158 | Hyd |
| | Optional | -4.912 | -7.449 | -2.158 | Hyd |
| | Optional | -8.047 | -6.301 | -6.811 | Hyd |
| x1083_0A-x1498_0A | Mandatory | -6.835 | -3.99 | -9.255 | Acc |
| | Optional | -2.403 | -7.764 | 1.32 | Hyd |
| | Optional | -5.963 | -8.561 | -1.504 | Hyd |
| | Optional | -8.047 | -6.301 | -6.811 | Hyd |
| x1140_0A-x1498_0A | Mandatory | -6.741 | -4.237 | -3.743 | Acc |
| | Mandatory | -6.835 | -3.99 | -9.255 | Acc |
| | Optional | -6.412 | -4.724 | -0.83 | Hyd |
| | Optional | -5.332 | -8.174 | -1.881 | Hyd |
| | Optional | -8.047 | -6.301 | -6.811 | Hyd |

Table 4: Pharmacophore features for docking merges against EV A71 2A protease

| Pair | Requirement | x | y | z | Type |
|---|---|---|---|---|---|
| x0310_0A-x0416_0A | Mandatory | 10.289 | 11.94 | 22.269 | Don |
| | Optional | 8.219 | 11.784 | 24.252 | Hyd |
| | Optional | 10.009 | 8.93 | 22.729 | Hyd |
| | Optional | 11.934 | 13.737 | 22.43 | Hyd |
| | Optional | 9.447 | 11.481 | 23.619 | Hyd |
| | Optional | 9.759 | 9.028 | 23.134 | Hyd |
| x0310_0A-x0556_0A | Mandatory | 10.289 | 11.94 | 22.269 | Don |
| | Mandatory | 6.244 | 9.398 | 26.831 | Don |
| | Optional | 8.219 | 11.784 | 24.252 | Hyd |
| | Optional | 10.009 | 8.93 | 22.729 | Hyd |
| | Optional | 11.934 | 13.737 | 22.43 | Hyd |
| | Optional | 8.695 | 11.508 | 23.717 | Hyd |
| | Optional | 4.995 | 8.454 | 28.88 | Hyd |
| | Optional | 4.995 | 8.454 | 28.88 | Hyd |
| x0416_0A-x0556_0A | Mandatory | 6.244 | 9.398 | 26.831 | Don |
| | Optional | 9.447 | 11.481 | 23.619 | Hyd |
| | Optional | 9.759 | 9.028 | 23.134 | Hyd |
| | Optional | 8.695 | 11.508 | 23.717 | Hyd |
| | Optional | 4.995 | 8.454 | 28.88 | Hyd |
| | Optional | 4.995 | 8.454 | 28.88 | Hyd |

Table 5: Pharmacophore features for docking merges against Zika NS2B

| Pair | Requirement | x | y | z | Type |
|---|---|---|---|---|---|
| x0089_0B-x1098_0B | Mandatory | -6.917 | 5.115 | -18.61 | Aro |
| | Mandatory | -4.785 | 5.51 | -20.319 | Don |
| | Mandatory | -9 | 4.657 | -17.727 | Aro |
| | Mandatory | -8.699 | 5.289 | -11.548 | Aro |
| | Mandatory | -9.161 | 4.117 | -14.146 | Don |
| | Optional | -2.549 | 4.447 | -19.752 | Hyd |
| | Optional | -9.427 | 4.563 | -18.786 | Hyd |
| | Optional | -2.549 | 4.447 | -19.752 | Hyd |
| | Optional | -7.384 | 5.028 | -17.559 | Hyd |
| | Optional | -9.345 | 4.386 | -19.035 | Hyd |
| | Optional | -7.729 | 5.164 | -17.427 | Hyd |
| | Optional | -8.825 | 4.899 | -12.652 | Hyd |
| | Optional | -7.729 | 5.164 | -17.427 | Hyd |
| x0429_0B-x1098_0B | Mandatory | -8.664 | 4.845 | -17.588 | Aro |
| | Mandatory | -9 | 4.657 | -17.727 | Aro |
| | Mandatory | -8.699 | 5.289 | -11.548 | Aro |
| | Mandatory | -9.161 | 4.117 | -14.146 | Don |
| | Optional | -9.99 | 4.459 | -17.653 | Hyd |
| | Optional | -8.059 | 4.976 | -16.345 | Hyd |
| | Optional | -9.345 | 4.386 | -19.035 | Hyd |
| | Optional | -7.729 | 5.164 | -17.427 | Hyd |
| | Optional | -8.825 | 4.899 | -12.652 | Hyd |
| | Optional | -7.729 | 5.164 | -17.427 | Hyd |
| x0687_0B-x0969_1B | Mandatory | -8.952 | 4.254 | -15.218 | Acc |
| | Mandatory | -6.504 | 5.241 | -18.609 | Aro |
| | Mandatory | -8.956 | 5.417 | -9.301 | Aro |
| | Mandatory | -9.077 | 1.514 | -9.134 | Don |
| | Optional | -7.121 | 5.464 | -17.383 | Hyd |
| | Optional | -9.356 | 4.727 | -18.369 | Hyd |
| | Optional | -7.121 | 5.464 | -17.383 | Hyd |
| | Optional | -8.708 | 5.605 | -11.972 | Hyd |

#### 7.5 Results of perfect versus bioisosteric merging using the Fragment Network

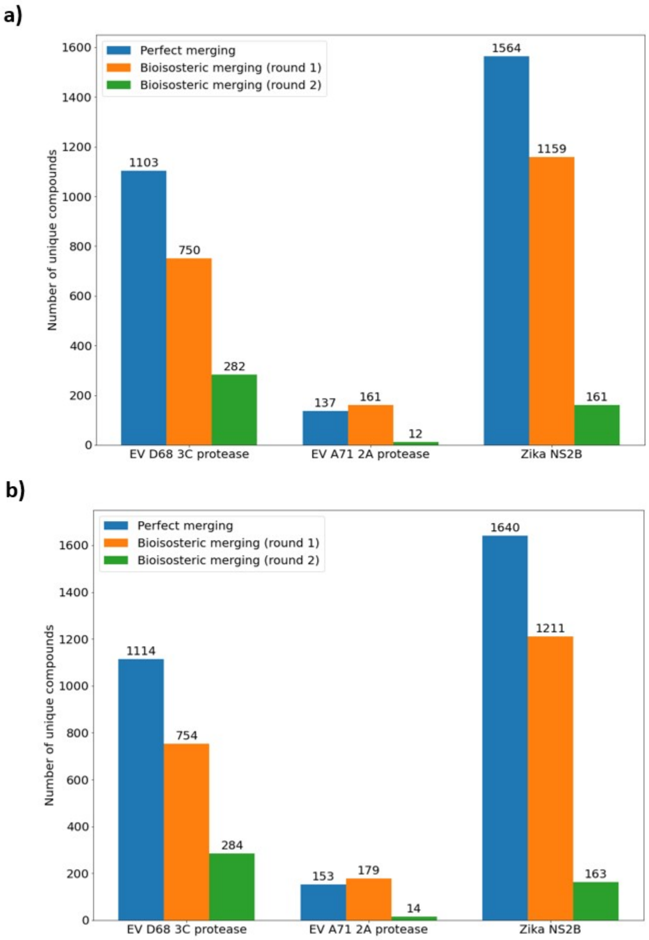


Figure 3: **Numbers of successfully filtered compounds from Fragment Network merging pipelines.** Compounds retrieved from the database by either perfect merging or bioisosteric merging, replacing one (round 1) or two (round 2) substructures are placed using an adapted Fragmenstein protocol [31] and minimized using PyRosetta [32]. The results are shown for compounds identified (**a**) before and (**b**) after performing an additional R-group search (to recapitulate substituents observed in the original fragments). Only the number of unique, successfully placed and filtered compounds with $SC_{RDKIT}$ values $\geq$0.55 are shown.

Table 6: Total hours for querying Fragment Network-derived compounds

| Target | EV D68 3C protease | EV A71 2A protease | Zika NS2B |
|---|---|---|---|
| Bioisosteric (round 1) | 186.31 | 1.96 | 419.85 |
| Perfect | 79.76 | 3.51 | 82.14 |
| Bioisosteric (round 2: strict search) | 0.41 | 0.39 | 0.23 |
| Bioisosteric (round 2: loose search) | 9.72 | 0.33 | 40.15 |
| R-group: bioisosteric (round 1) | 0.02 | 0.01 | 0.01 |
| R-group: perfect | 0.02 | 0 | 0.02 |
| R-group: bioisosteric (round 2: strict search) | 0 | 0 | 0 |
| R-group: Bioisosteric (round 2: loose search) | 0.01 | 0 | 0 |
| Total time | 276.25 | 6.2 | 542.4 |
| Estimated average time per fragment pair* | 3.63 | 1.13 | 6.42 |

*Figures show the total times (in hours) for running each individual query. Up to 8 queries were run in parallel on a VM with 120 CPUs. *Calculated based on asymmetric fragment pairs (i.e. a pair may be repeated). EV, enterovirus.*

Table 7: Total CPU hours for filtering Fragment Network-derived compounds

| | EV D68 3C protease | EV A71 2A protease | Zika NS2B |
|---|---|---|---|
| Bioisosteric (round 1) | 1754.81 | 87.8 | 791.9 |
| Perfect | 2492.74 | 138.74 | 1184.67 |
| Bioisosteric (round 2) | 1149.65 | 24.26 | 115.66 |
| R-group: bioisosteric (round 1) | 30.2 | 6.5 | 23.66 |
| R-group: perfect | 44.7 | 6.04 | 45.16 |
| R-group: bioisosteric (round 2: loose search) | 13.67 | 0.48 | 5.18 |
| R-group: Bioisosteric (round 2: strict search) | 9.82 | 0.98 | 3.37 |
| Total time | 5495.59 | 264.8 | 2169.6 |
| Estimated average time per fragment pair* | 72.3 | 48.1 | 25.7 |

**Calculated based on asymmetric fragment pairs (i.e. a pair may be repeated). EV, enterovirus*

Table 8: Numbers of molecules passing through the merging pipeline

| Target | Method | N molecules from database | N molecules entering minimization | N placed molecules | N filtered molecules | N filtered molecules (after R-group exp) |
|---|---|---|---|---|---|---|
| EV D68 3C protease | P | 129,328 (65,753) | 129,328 (65,753) | 30,063 (18,387) | 1,849 (1,103) | 1,860 (1,114) |
| | B1 | 217,411 (99,245) | 188,599 (91,139) | 27,865 (15,532) | 1,351 (750) | 1,355 (754) |
| | B2 | 60,569 (14,577) | 60,569 (14,577) | 18,927 (4,709) | 900 (282) | 903 (284) |
| EV A71 2A protease | P | 22,747 (13,351) | 22,747 (13,351) | 1,555 (1,360) | 156 (137) | 178 (153) |
| | B1 | 32,297 (20,106) | 26,123 (16,245) | 1,440 (1,104) | 193 (161) | 221 (179) |
| | B2 | 2,631 (1,836) | 2,631 (1,836) | 350 (248) | 13 (12) | 15 (14) |
| Zika NS2B | P | 145,401 (74,263) | 145,401 (74,263) | 30,349 (17,150) | 2,953 (1,564) | 3,050 (1,640) |
| | B1 | 171,528 (71,431) | 156,658 (61,450) | 24,075 (11,601) | 2,039 (1,159) | 2,108 (1,211) |
| | B2 | 12,690 (4,898) | 12,690 (4,898) | 5,135 (1,702) | 674 (161) | 677 (163) |

*Brackets indicate the numbers of unique molecules. B1, first round of bioisosteric merging; B2, second round of bioisosteric merging; EV, enterovirus; P, perfect merging.*

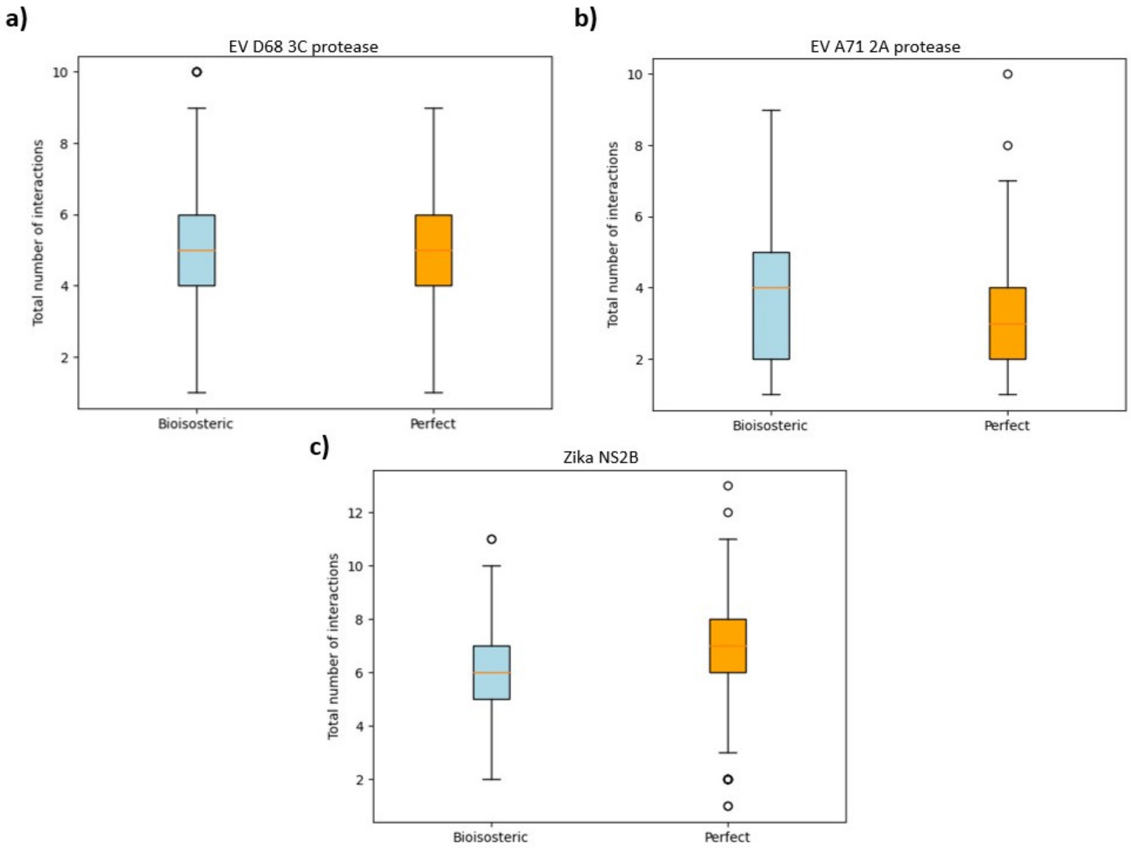


Figure 4: **Total number of predicted interactions made by each merge.** The total number of predicted interactions for each compound proposed by bioisosteric merging and perfect merging for (**a**) enterovirus (EV) D68 3C protease, (**b**) EV A71 2A protease and (**c**) Zika NS2 protease. Interactions are calculated using ProLIF. Only compounds with a $SC_{RDKIT}$ score $\geq$ 0.55 and represent 'true merges' (potentially replicating an interaction from each fragment) are shown. The results are shown to include all compounds after R-group expansions.

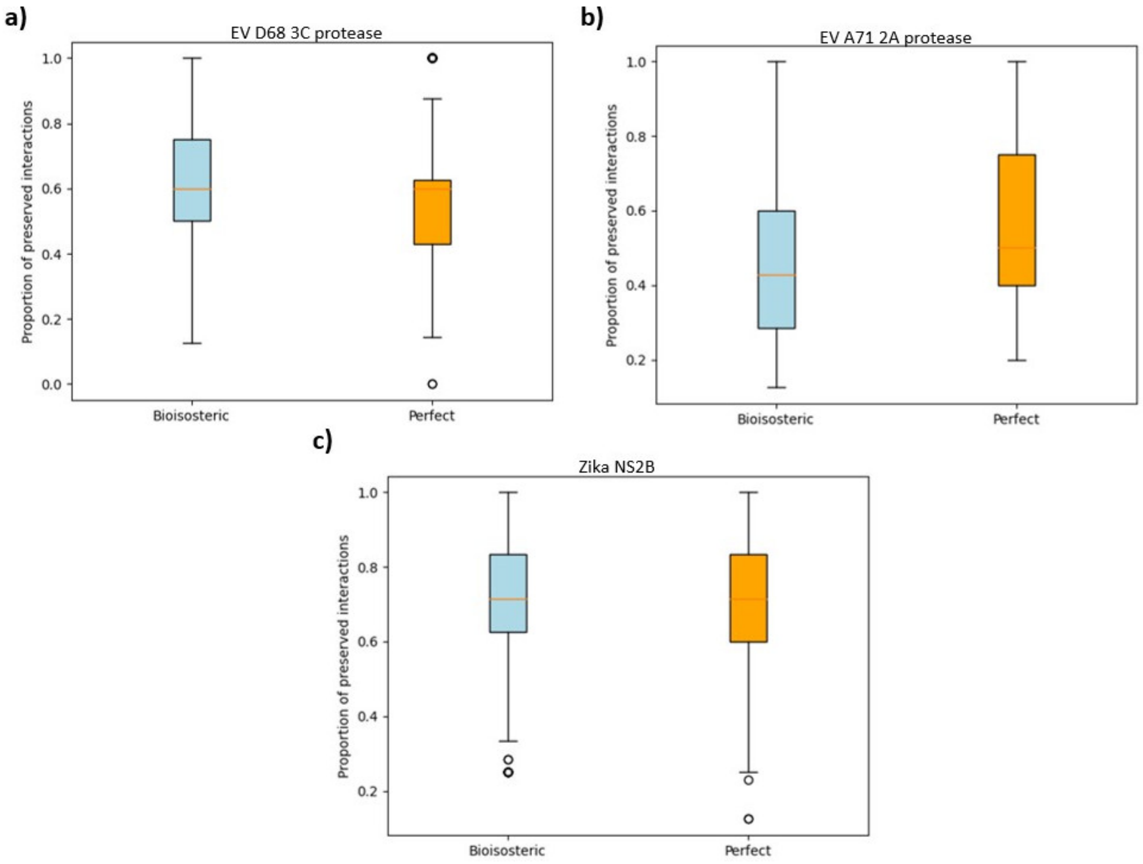


Figure 5: **Fraction of preserved interactions.** The proportion of preserved predicted interactions observed at the residue-level (that is, unique interactions that are seen in the parent fragments that are replicated by the merge) for each compound proposed by bioisosteric merging and perfect merging for (**a**) enterovirus (EV) D68 3C protease, (**b**) EV A71 2A protease and (**c**) Zika NS2B. Interactions are calculated using ProLIF. Only compounds with a $SC_{\mathrm{RDKIT}}$ score $\geq 0.55$ and represent 'true merges' (replicating an interaction from each fragment) are shown. The results are shown to include all compounds after R-group expansions.

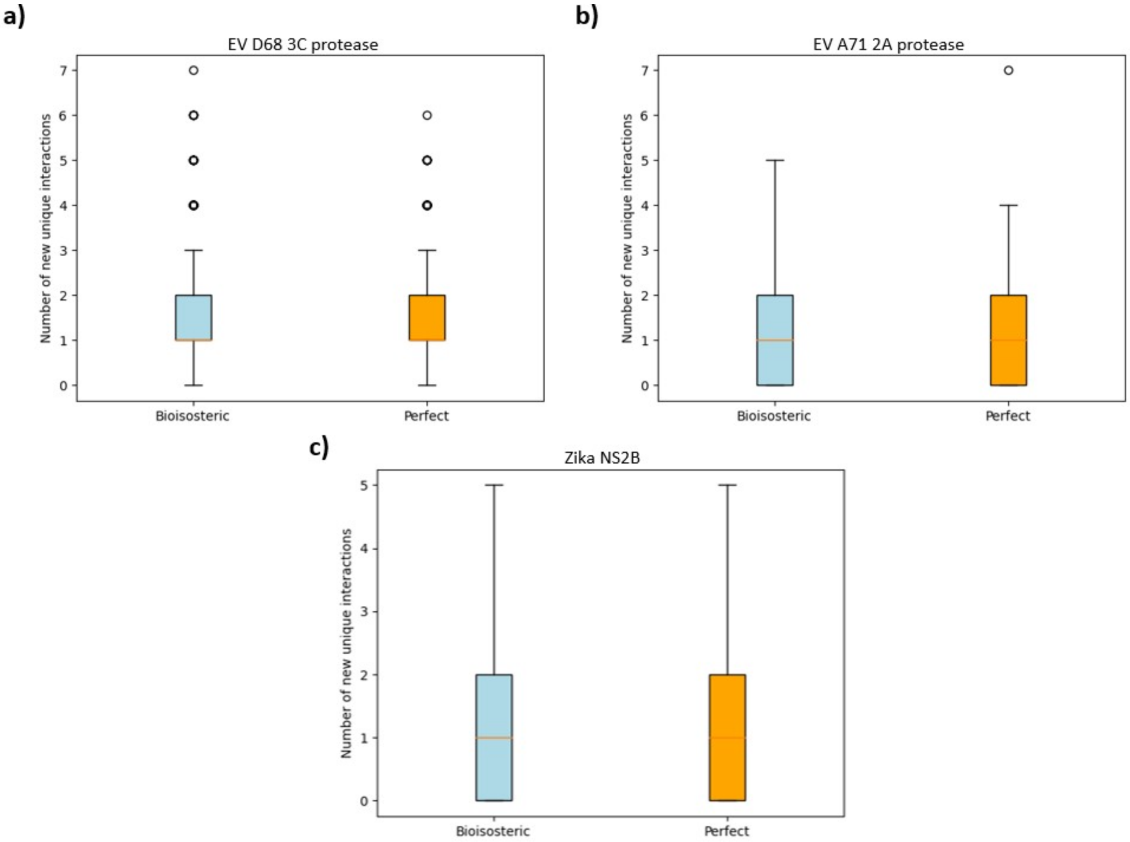


Figure 6: **Total number of predicted new unique interactions.** The total number of new unique predicted interactions observed at the residue-level (that is, interactions that are not seen in the parent fragments) for each compound merge proposed by bioisosteric merging and perfect merging for (**a**) enterovirus (EV) D68 3C protease, (**b**) EV A71 2A protease and (**c**) Zika NS2B. Interactions are calculated using ProLIF. Only compounds with a $SC_{RDKIT}$ score $\geq$ 0.55 and represent 'true merges' (replicating an interaction from each fragment) are shown. The results are shown to include all compounds after R-group expansions.

Table 9: Clustering analysis for bioisosteric and perfect merging pipelines.

| Distance threshold | Cluster composition | EV D68 3C protease | EV A71 2A protease | Zika NS2B |
|---|---|---|---|---|
| 0.3 | P + B | 39 | 7 | 92 |
| | P | 801 | 122 | 1,226 |
| | B | 775 | 133 | 1,019 |
| 0.4 | P + B | 52 | 9 | 129 |
| | P | 547 | 85 | 735 |
| | B | 551 | 96 | 644 |

*B, bioisosteric merging; EV, enterovirus; P, perfect merging.*

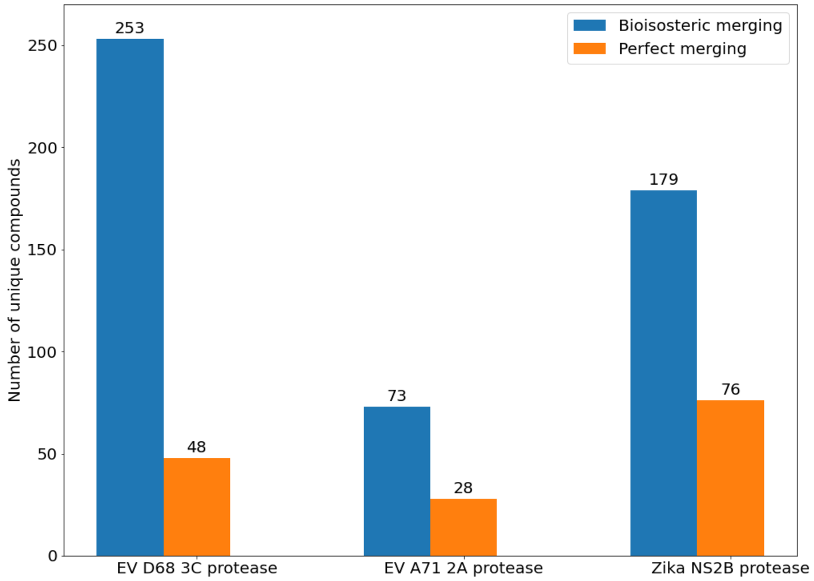


Figure 7: **Bioisosteric merging increases the numbers of unique substructures represented in follow-up compounds.** The numbers of unique substructures contributing to the final sets of merge compounds are shown across all targets. Bioisosteric merging increases the numbers of substructures observed compared with perfect merging. EV, enterovirus.

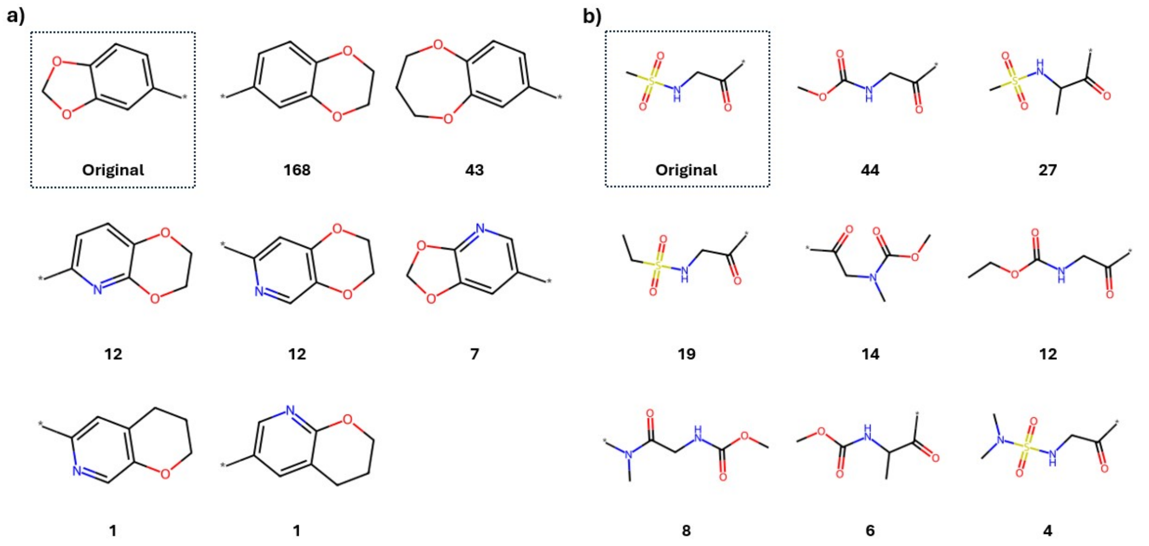


Figure 8: **Example substructure replacements from bioisosteric merging.** Example substructure replacements seen for two susbtructures used to form merges from the enterovirus D68 3C protease fragment screen. An example for a substructure containing (**a**) fused rings and (**b**) no rings are shown, together with the original substructures. Numbers indicate the number of compounds that contain the replacement substructure. The results are shown to include all compounds after R-group expansions.

#### 7.6 Comparison of Fragment Network vs RDock

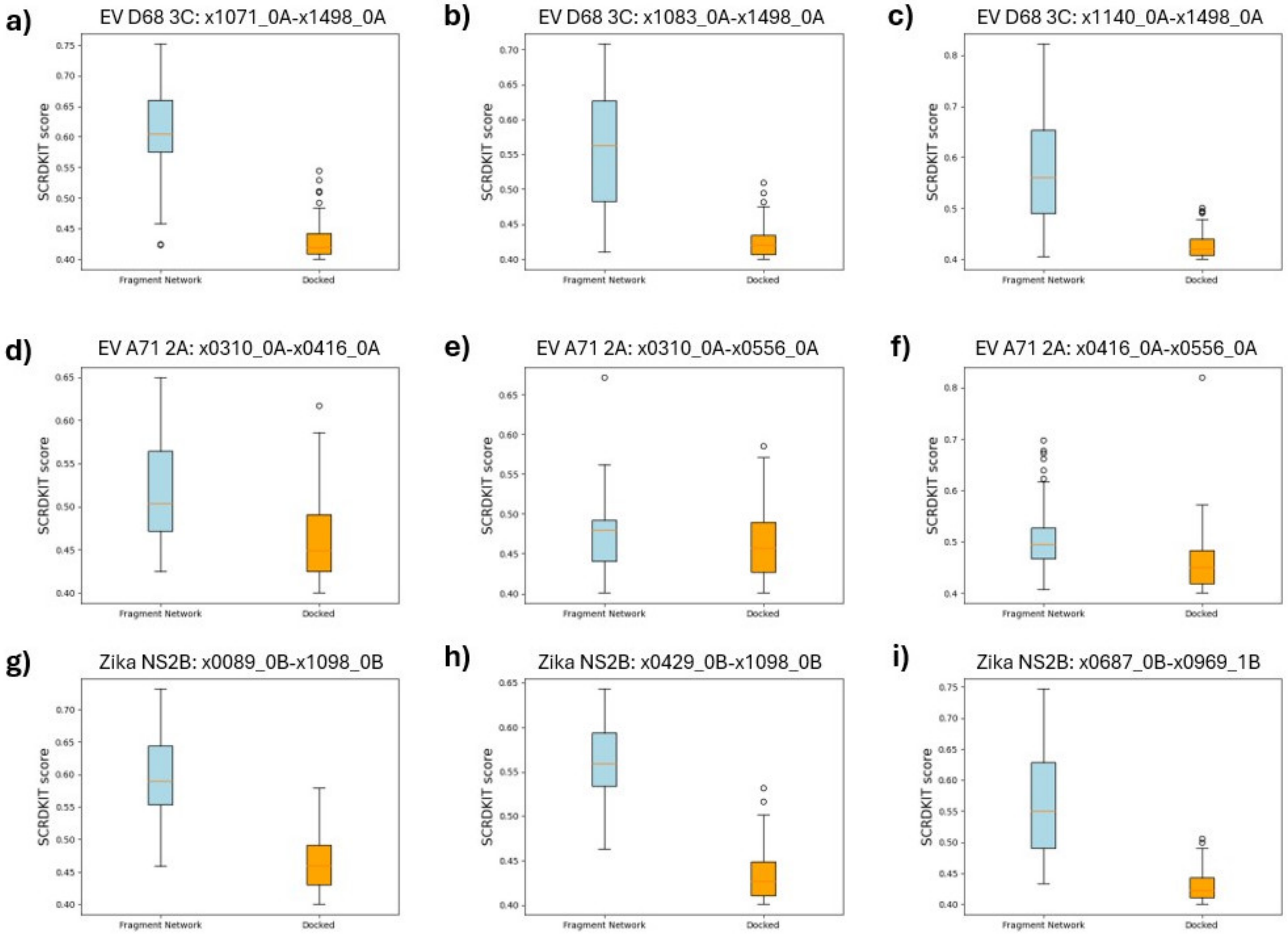


Figure 9: **$SC_{RDKIT}$ scores made by top 100 compounds identified using the Fragment Network merging pipelines versus pharmacophore-constrained docking.** EV, enterovirus.

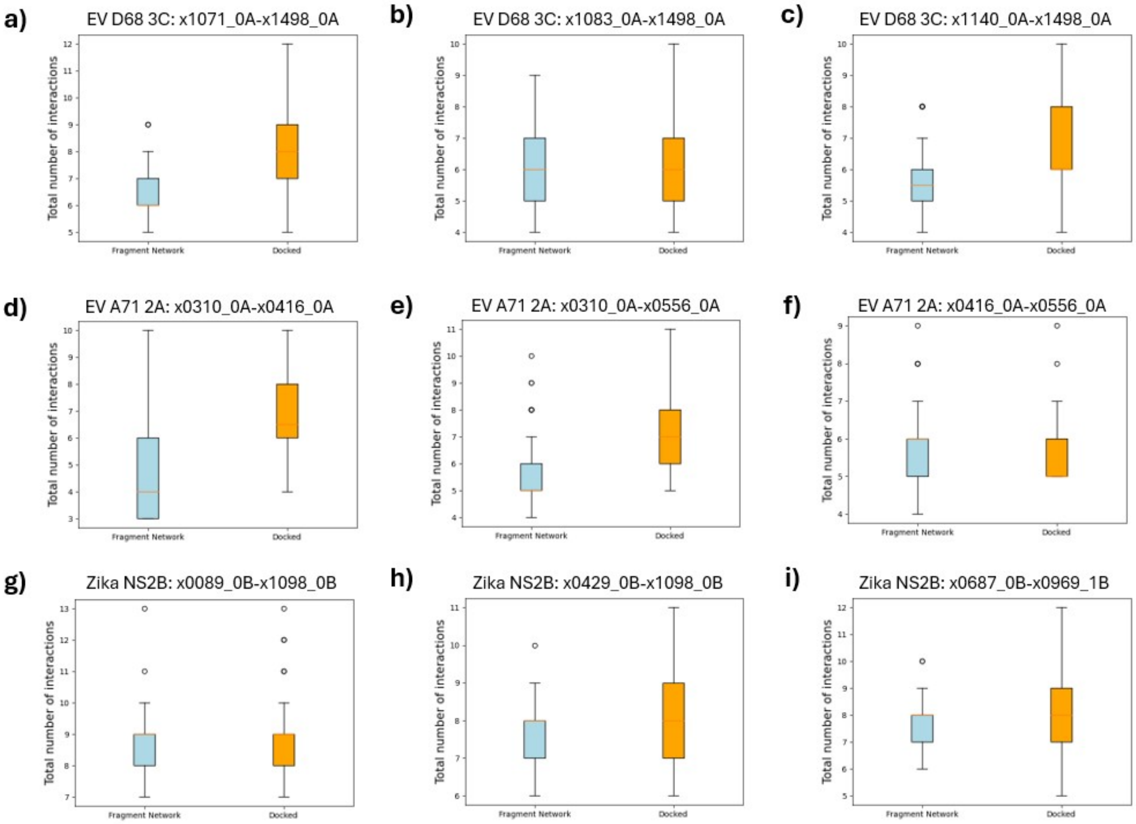


Figure 10: **Total number of predicted interactions made by top 100 compounds identified using the Fragment Network merging pipelines versus pharmacophore-constrained docking.** EV, enterovirus.

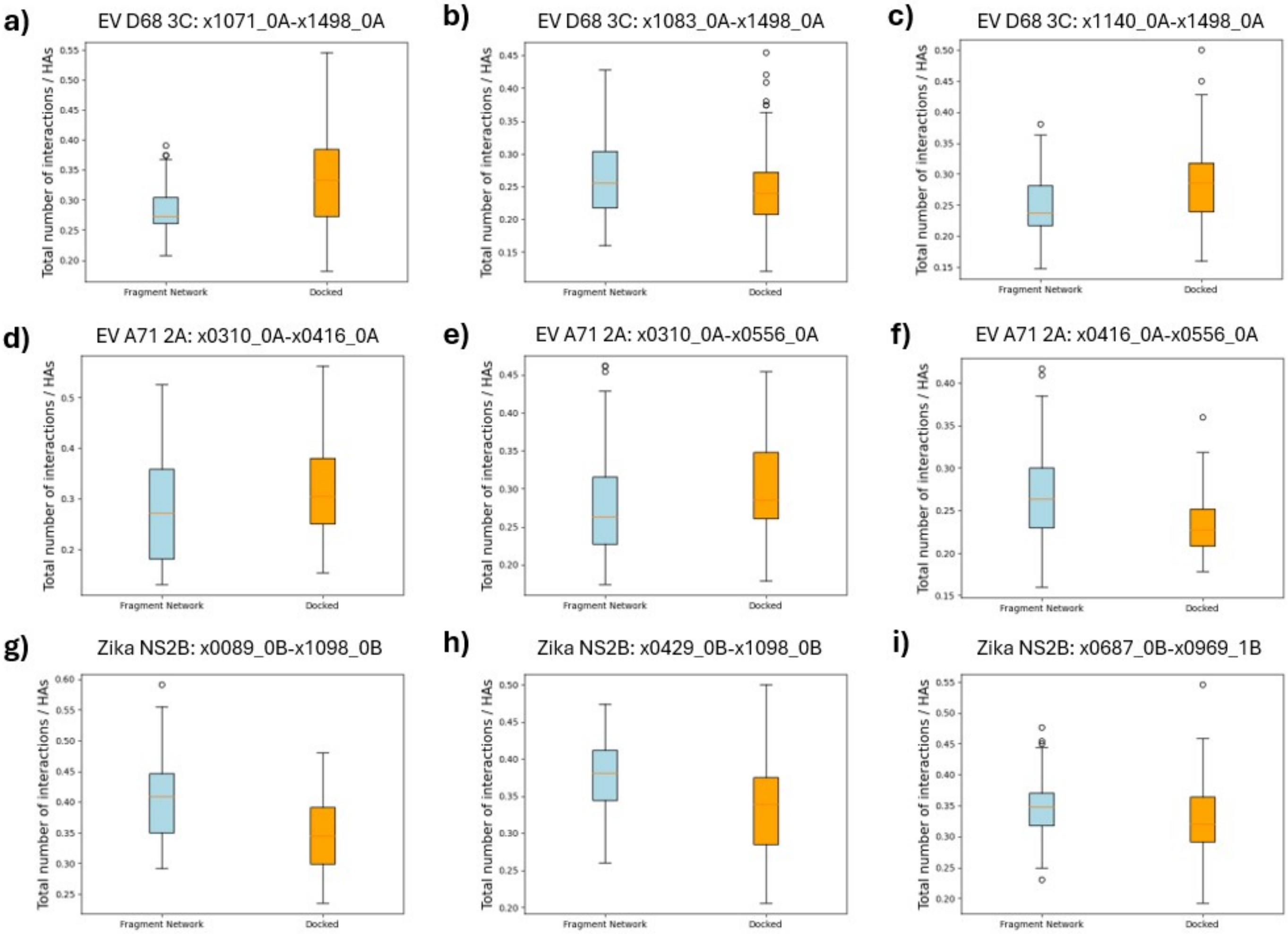


Figure 11: **Number of predicted interactions made (normalized by heavy atom count) made by top 100 compounds identified using the Fragment Network merging pipelines versus pharmacophore-constrained docking.** EV, enterovirus.

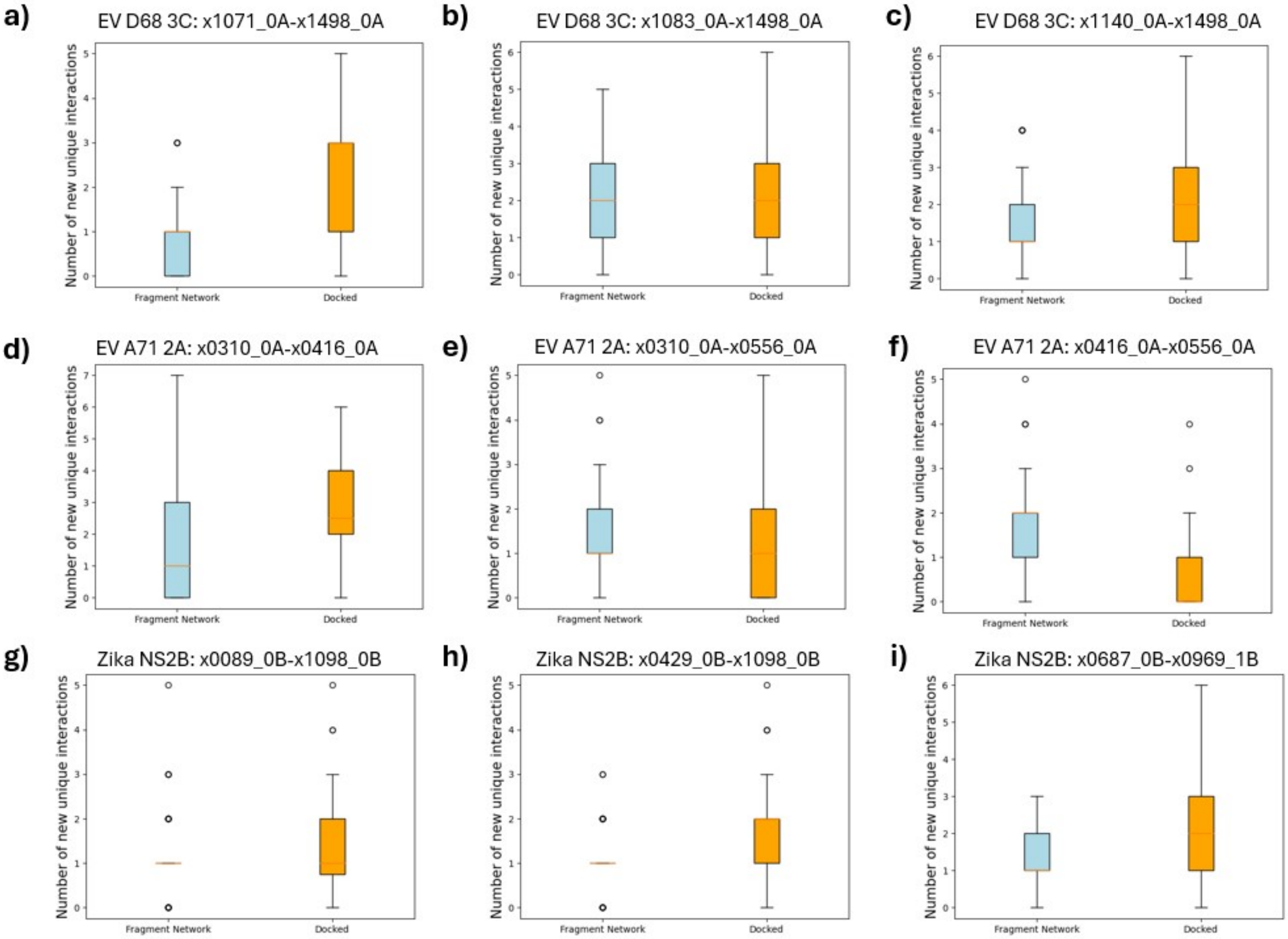


Figure 12: **Number of new predicted interactions not seen in parent fragments made by top 100 compounds identified using the Fragment Network merging pipelines versus pharmacophore-constrained docking.** EV, enterovirus.

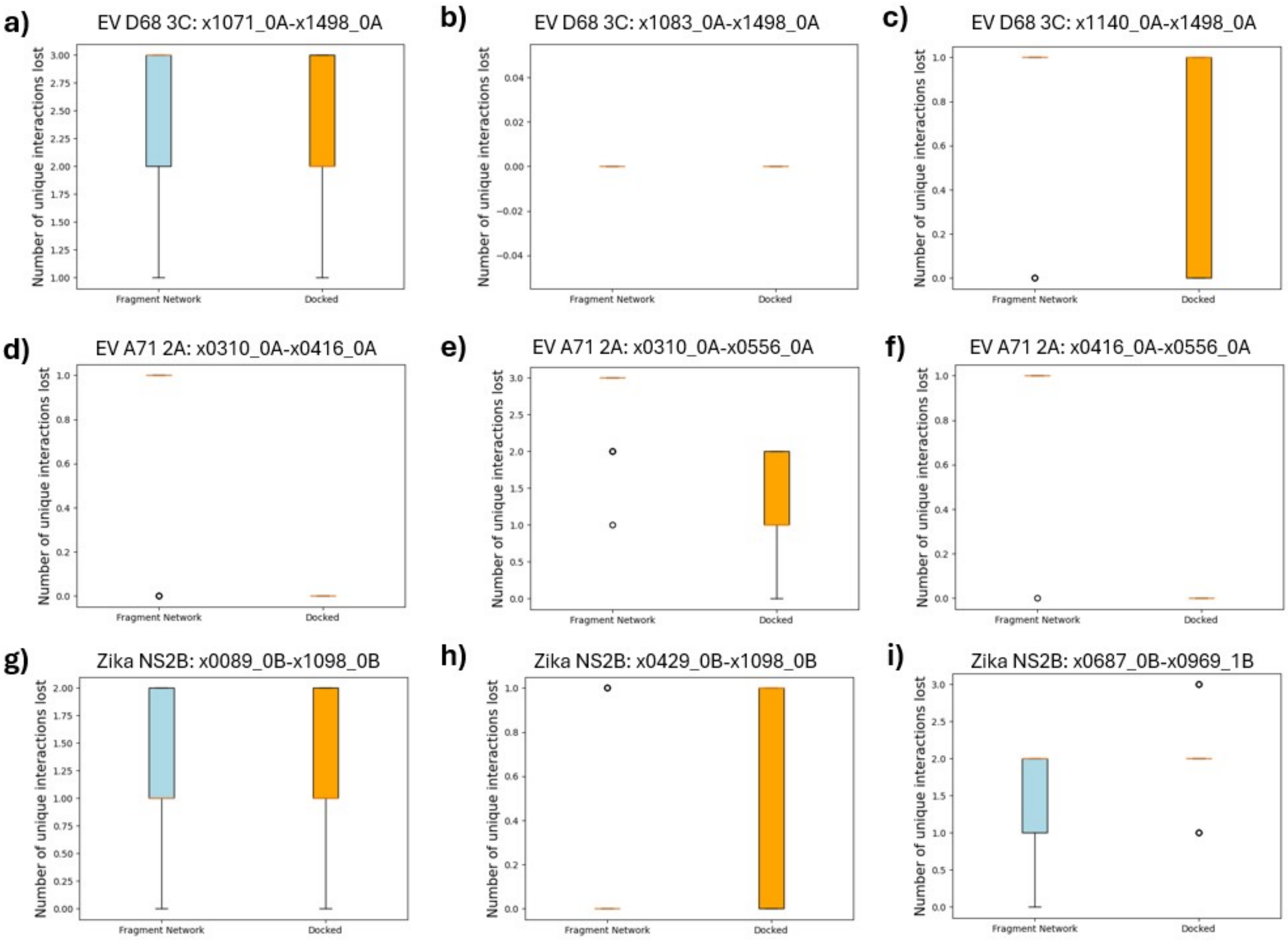


Figure 13: **Number of unique interactions made by parent fragments that are lost for the top 100 compounds identified using the Fragment Network merging pipelines versus pharmacophore-constrained docking.** The number of unique interactions made by the parent fragments for each merge that are predicted to be lost by the proposed merge. EV, enterovirus.

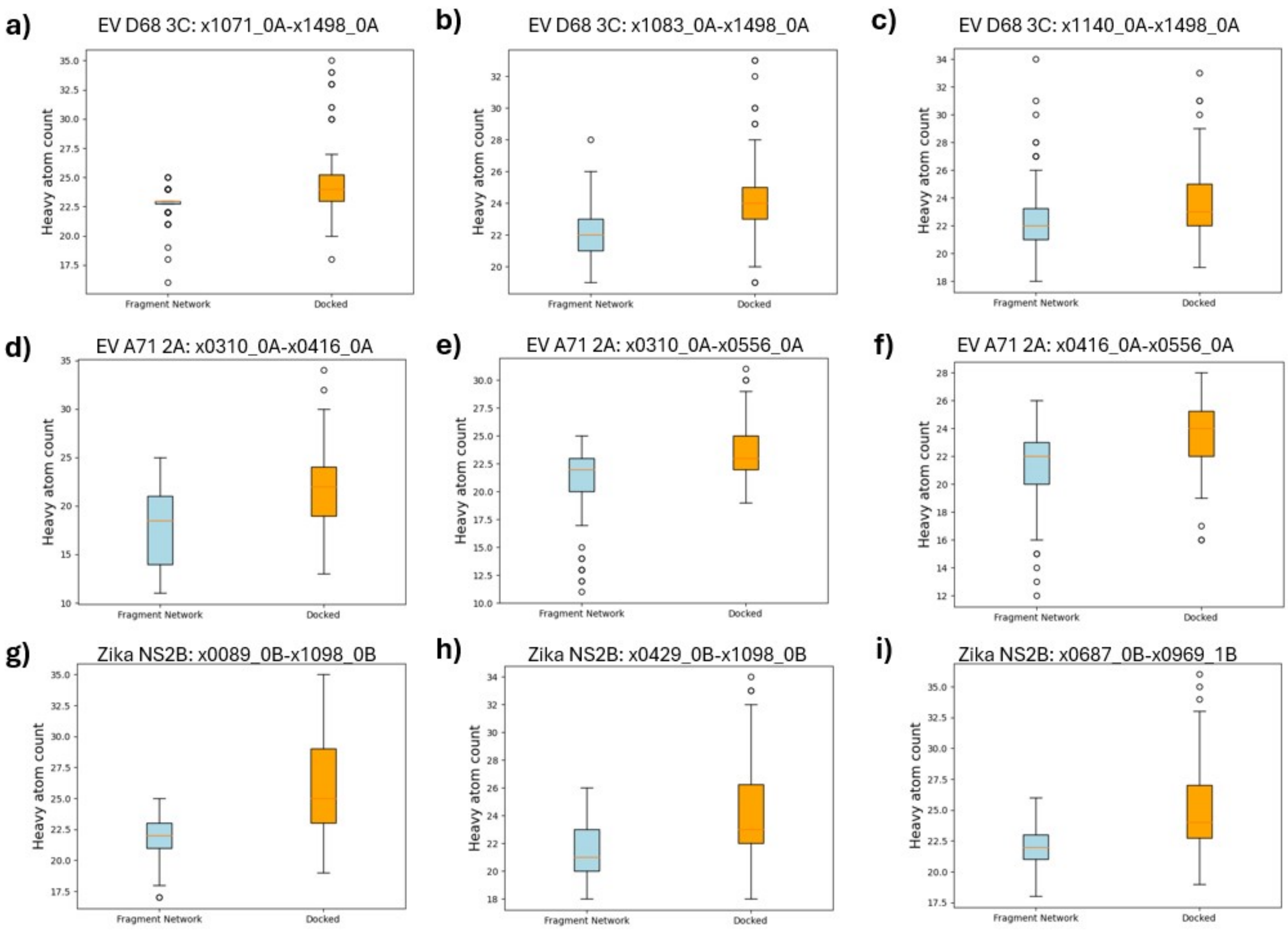


Figure 14: **Heavy atom count for top 100 compounds identified using the Fragment Network merging pipelines versus pharmacophore-constrained docking.** EV, enterovirus.

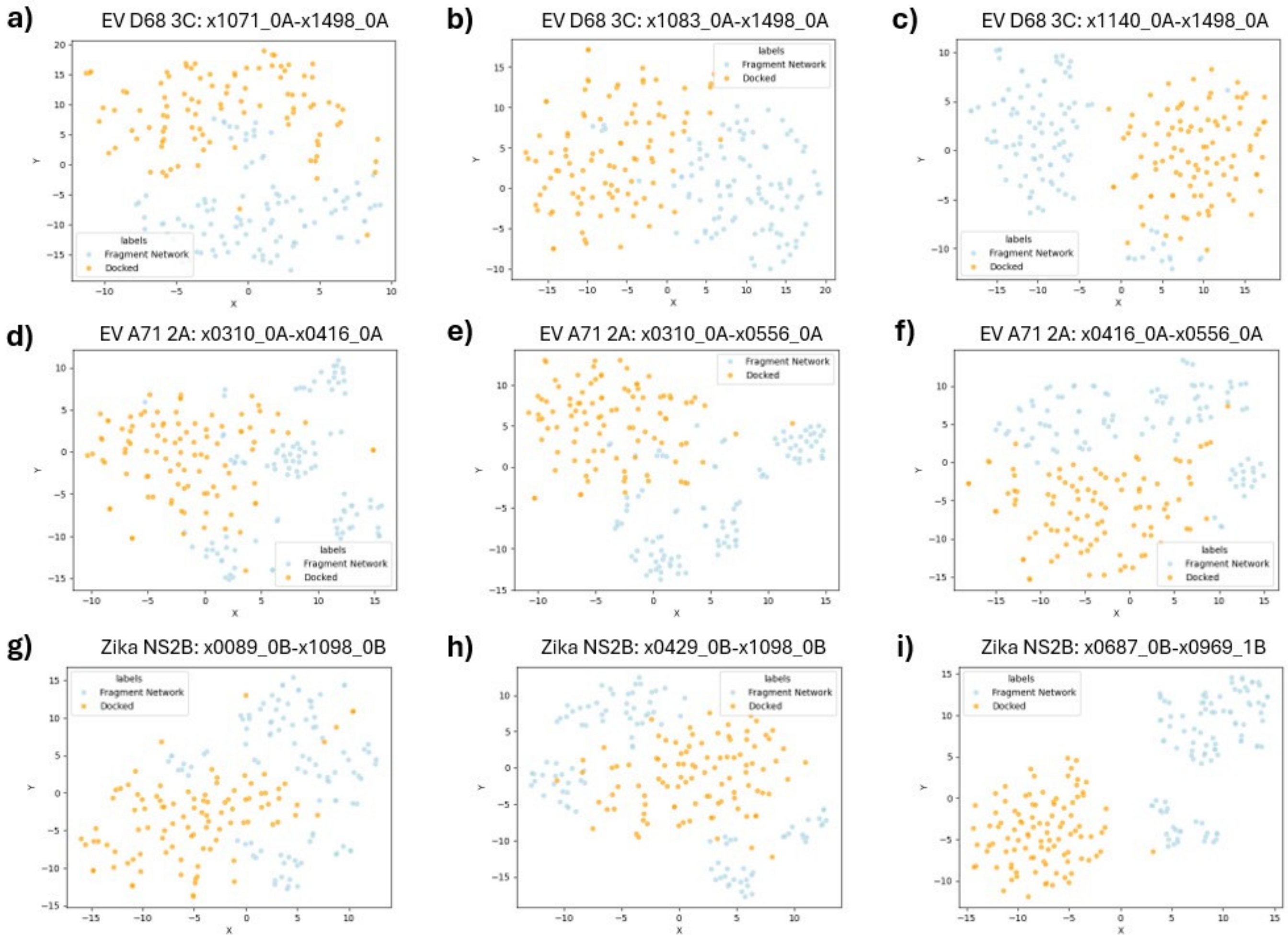


Figure 15: **TSNEs for top 100 compounds identified using the Fragment Network merging pipelines versus pharmacophore-constrained docking.** EV, enterovirus.

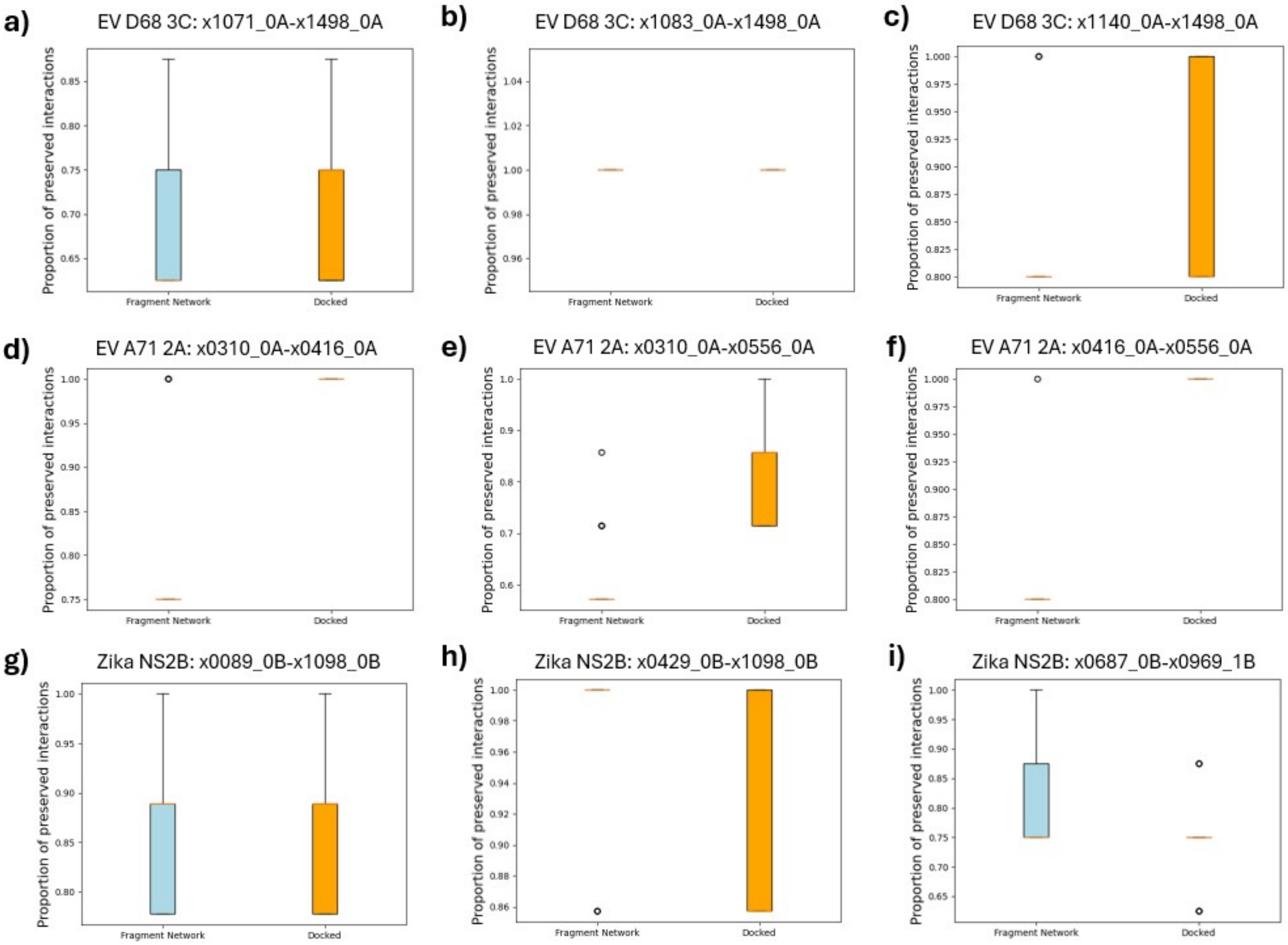


Figure 16: **The fraction of preserved interactions from the parent fragments for top 100 compounds identified using the Fragment Network merging pipelines versus pharmacophore-constrained docking.** EV, enterovirus.

Table 10: The number of clusters containing the top-ranked Fragment Network or rDock-derived compounds.

| Target | Pair | Number of FN clusters | Number of rDock clusters |
|---|---|---|---|
| EV D68 3C protease | x1071_0A-x1498_0A | 56 | 99 |
| | x1083_0A-x1498_0A | 73 | 100 |
| | x1140_0A-x1498_0A | 70 | 100 |
| EV A71 2A protease | x0310_0A-x0416_0A | 62 | 100 |
| | x0310_0A-x0556_0A | 60 | 99 |
| | x0416_0A-x0556_0A | 65 | 99 |
| Zika NS2B | x0089_0B-x1098_0B | 58 | 97 |
| | x0429_0B-x1098_0B | 60 | 99 |
| | x0687_0B-x0969_1B | 54 | 100 |

*Clustered using the Taylor-Butina algorithm using Tanimoto similarity (threshold of 0.4) and Morgan fingerprints (radius 2; 1,024 bits). EV, enterovirus; FN, Fragment Network.*

#### 7.7 Retrospective analysis

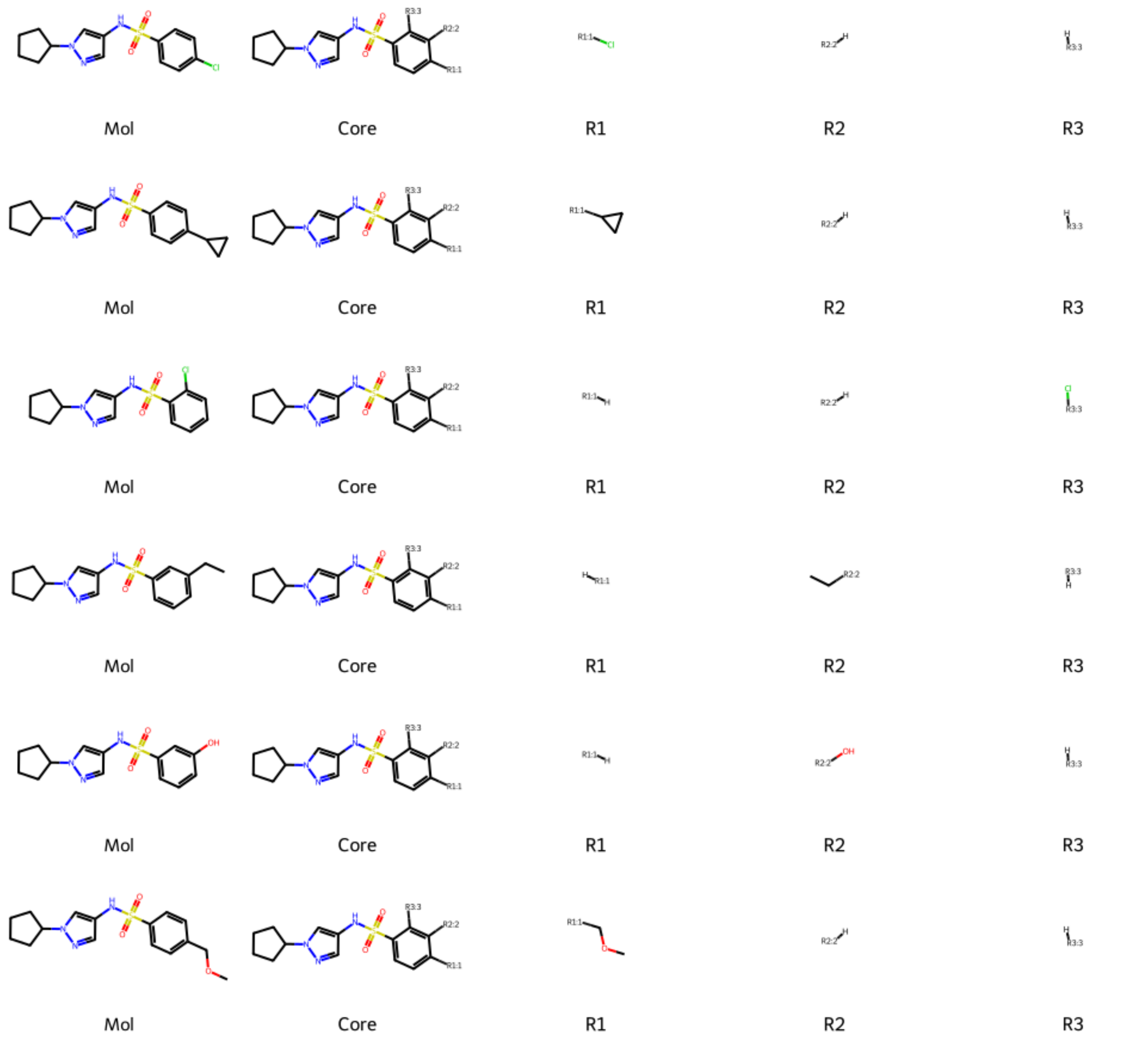


Figure 17: **R-group decomposition for molecules similar to substituent 3d against WDR5–MYC.** Fragment Network-identified merges of fragments that bind to the WDR5–MYC complex. The identified merges incorporate the pyrazole ring identified for substituent 3d (manually designed in [35]) and also various substituents, which were outlined in the displayed R-group decomposition.

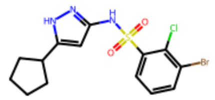

Mol

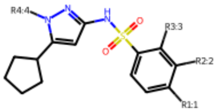

Core

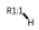

R1



R2

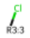

R3

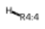

R4

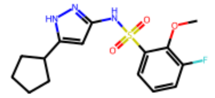

Mol

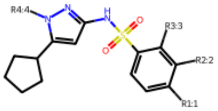

Core



R1

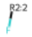

R2

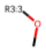

R3

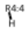

R4

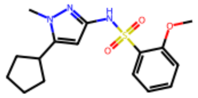

Mol

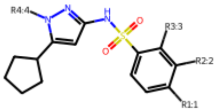

Core



R1

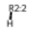

R2

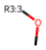

R3

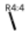

R4

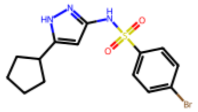

Mol

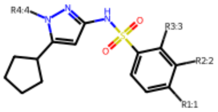

Core

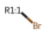

R1



R2

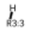

R3



R4

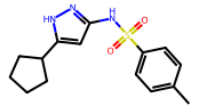

Mol

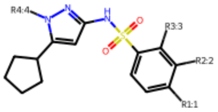

Core

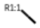

R1



R2

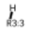

R3



R4

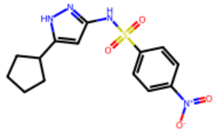

Mol

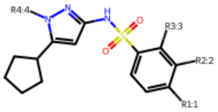

Core

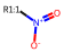

R1

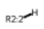

R2

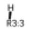

R3



R4

[h]

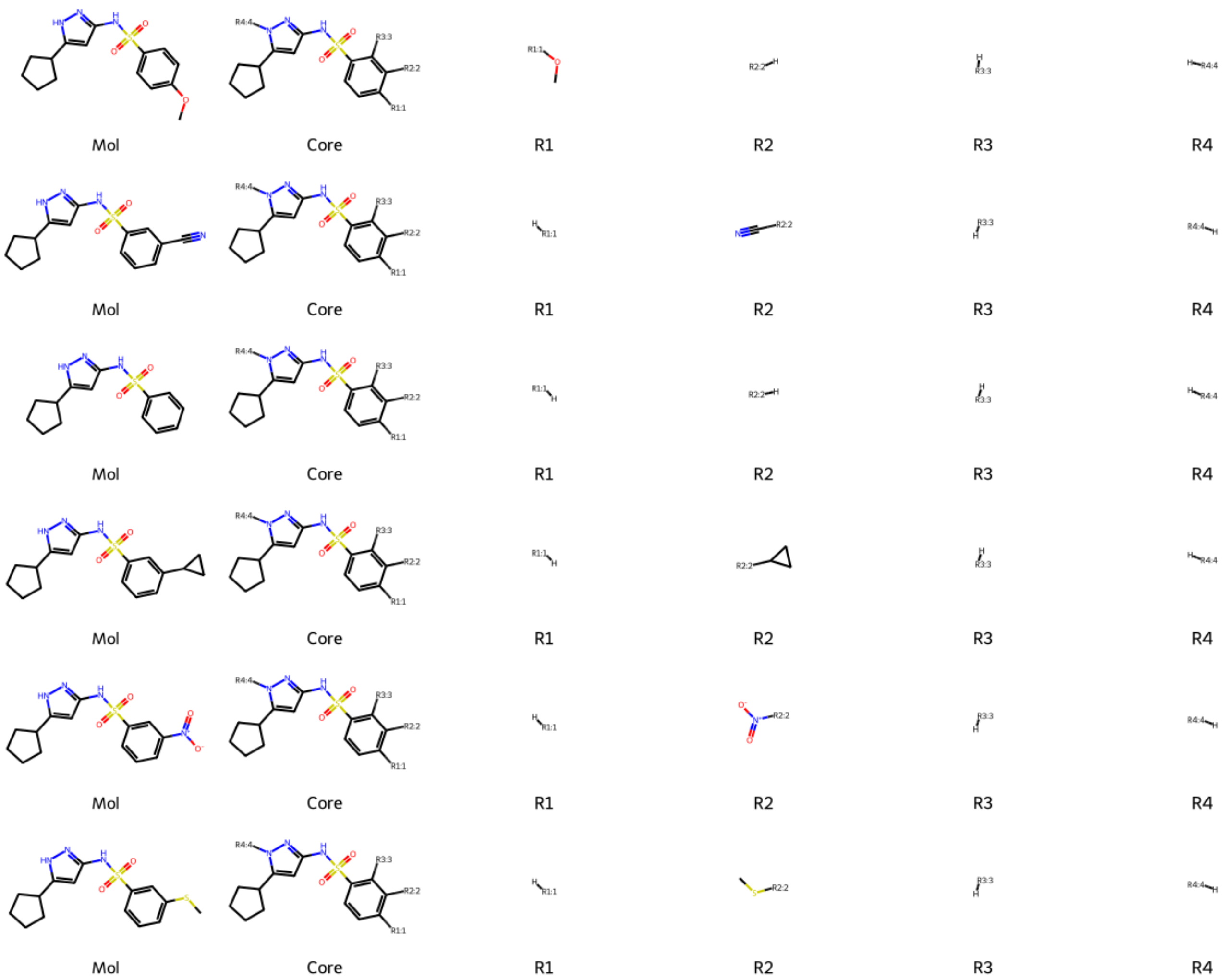


Figure 18: **R-group decomposition for molecules similar to substituent 3b against WDR5–MYC.** Fragment Network-identified merges of fragments that bind to the WDR5–MYC complex. The identified merges incorporate the pyrazole ring identified for substituent 3b (manually designed in [35]) and also various substituents, which were outlined in the displayed R-group decomposition.
